# Beyond pulsed inhibition: Alpha oscillations modulate attenuation and amplification of neural activity in the awake resting-state

**DOI:** 10.1101/2022.03.03.482657

**Authors:** Fabrizio Lombardi, Hans J. Herrmann, Liborio Parrino, Dietmar Plenz, Silvia Scarpetta, Anna Elisabetta Vaudano, Lucilla de Arcangelis, Oren Shriki

## Abstract

The alpha rhythm is a distinctive feature of the awake resting-state of the human brain. Recent evidence suggests that alpha plays an active role in information processing, modulating behavioral and cognitive performance. However, the functional role of alpha oscillations in the resting-state neuronal dynamics remains poorly understood. To address this question, we investigate collective neural activity during resting wake and NREM sleep, a physiologic state with marginal presence of alpha rhythm. We show that, during resting wake, alpha oscillations drive an alternation of attenuation and amplification bouts in neural activity. Our analysis indicates that inhibition is activated in pulses that last a single alpha cycle and gradually suppress neural activity, while excitation is successively enhanced over timescales of a few alpha cycles to amplify neural activity. Furthermore, we show that long-term, intermittent fluctuations in alpha amplitude—known as the “waxing and waning” phenomenon—are associated with an attenuation-amplification mechanism acting over the timescales of several seconds and described by a power law decay of the activity rate in the “waning” phase. Importantly, we do not observe such dynamics during NREM sleep. The results suggest that the alpha rhythm acts as a “pacemaker” for the alternation of inhibition and excitation bouts across multiple timescales, the “waxing and waning” being a long-term control mechanism of cortical excitability. The amplification regime observed beyond the timescales of the individual alpha cycle suggests in turn that alpha oscillations might modulate the intensity of neural activity not only through pulses of inhibition, as proposed in the pulsed inhibition hypothesis, but also by timely enhancing excitation (or dis-inhibition).

## Introduction

The mammalian brain exhibits complex rhythmic dynamics that span a broad range of frequencies [1]. Brain rhythms emerge as periodic amplitude fluctuations in electro-physiological recordings, which result from the synchronous activation of large populations of neurons [2, 3]. Oscillations in distinct frequency bands have been associated with different brain functions and physiological states [2, 4, 5]. Among these brain rhythms, oscillations in the alpha band (8-13 Hz) play a prominent role in human brain activity. Characteristic of the eyes-closed, awake restingstate, alpha oscillations have been associated with the processes of task disengagement [6, 7], perceptual learning and suppression of visual activity [8, 9], facilitation of periodic sampling of visual information, and, more generally, with propagation of activity throughout the brain, modulation of communication between brain regions, and feedback processing within and across brain regions [10, 11, 12, 13]. Indeed, a number of studies indicate that changes in cognitive performance are accompanied by rhythmic modulation of alpha amplitude and power, suggesting in particular an inverse relationship between alpha activity and neural firing [6, 8, 14, 15]. Amplitude fluctuations occur across multiple timescales and show non-trivial features, such as long-range temporal correlations (LRTC) [16, 17] and non-Gaussian bimodal distributions of power [18], which suggests the existence of two distinct modes—higher- and lower-power modes [18]. In particular, periodic fluctuations in amplitude, known as a “waxing and waning”, have long been considered a key feature of alpha oscillations [19, 20].

Several models have been proposed to explain the emergence of alpha oscillations and their dynamic characteristics, from mutually coupled excitatory (E) and inhibitory (I) spiking and stochastic neurons [21, 22], to adaptive neural networks [23] and more realistic thalamic and corticothalamic mechanistic models [24, 25, 26, 27]. In particular, taking into account corticothalamic loop mechanisms, the neural field model by Freyer et al. [26] reproduces spontaneous jumps between lower- and higher-power alpha modes in the presence of a subcritical Hopf bifurcation. Apart from the awake resting-state, alpha oscillations can also be marginally observed in other physiologic states, such as rapid eye movement (REM) sleep and stage 1 of non-rapid eye movement (NREM) sleep, or during EEG arousals that may occur in REM and NREM sleep [4, 28, 29].

Despite the advances in understanding the generation of the alpha rhythm and its influence on behavioral performance, the functional role of alpha oscillations in brain dynamics remains to a large extent not understood. Recent studies suggest that alpha oscillations mediate cortical inhibition. However, the nature of this inhibition, as well as its effects on collective neural dynamics, is still not known. One of the hypotheses that has recently gained consensus is that alpha-mediated inhibition is delivered in rhythmic pulses [30, 31, 8, 13], a mechanism that could allow for selective information processing during the periods of relative excitation [32].

Verifying directly this hypothesis requires complex, simultaneous multiscale recordings of neural activity. Yet, we argue that it should be possible to identify the hallmarks of alpha-mediated, pulsed inhibition through the analysis of neural dynamics that are readily accessible. To this end, we note that, within the pulsed inhibition hypothesis, alternating states of inhibition and excitation should result in periods of attenuation and enhancement of collective neural activity rhythmically modulated by alpha oscillations. To verify this assumption and, at the same time, to further clarify the functional role of the alpha rhythm in collective neural dynamics, we study neural activity cascades during resting wake and NREM sleep, where alpha oscillations are only marginally present [4, 29]. Neural activity cascades, termed neuronal avalanches, are spatio-temporal patterns of activity with no characteristic size, time, or spatial scale [33, 34, 35, 36, 37], which coexist with neural oscillations [38, 39, 40, 17, 41, 23]. By dissecting the dynamics of neuronal avalanches in relation to the alpha rhythm, we show that alpha oscillations modulate bouts of neural activity attenuation and amplification unfolding over multiple timescales—from hundreds of milliseconds to seconds. This attenuation-amplification mechanism is not present during NREM sleep, and suggest that the alpha rhythm mediates the timing of both inhibition and excitation periods in the awake resting-state. Importantly, our analysis provides a first quantitative description of the collective neural dynamics underlying the “waxing and waning” of the alpha rhythm [19, 42, 18]. We show that the “waxing and waning” is a long-term mechanism for the regulation of resting-state network excitability, which is intermittent rather than periodic in nature. The results suggest that, in the awake resting state, alpha oscillations modulate the intensity of neural activity not only through pulses of inhibition, as in the pulsed inhibition hypothesis, but also by timely enhancing excitation (or dis-inhibition), an effect that was not previously described.

## Results

### Dynamics of neural activity cascades during the awake resting-state

To characterize collective brain dynamics in relation to the alpha rhythm, we analyzed MEG and EEG recordings of the awake, eyes-closed, resting-state brain activity (STAR Methods). First, we proceeded to identify spatiotemporal cascades of neural activity across the sensor arrays. To this end, we mapped each continuous, broadband sensor signal into a sequence of discrete events. These are defined as the extremes of large positive and negative signal deflections exceeding an amplitude threshold *h* (Fig. 1A). In practice, for each excursion beyond the threshold, a single event is identified at the most extreme value (maximum for positive excursions and minimum for negative excursions; blue dots in Fig. 1A) (STAR Methods). This procedure preserves most of the collective features encoded in the continuous signals across the sensor array [43, 36, 37].

**Figure 1.**
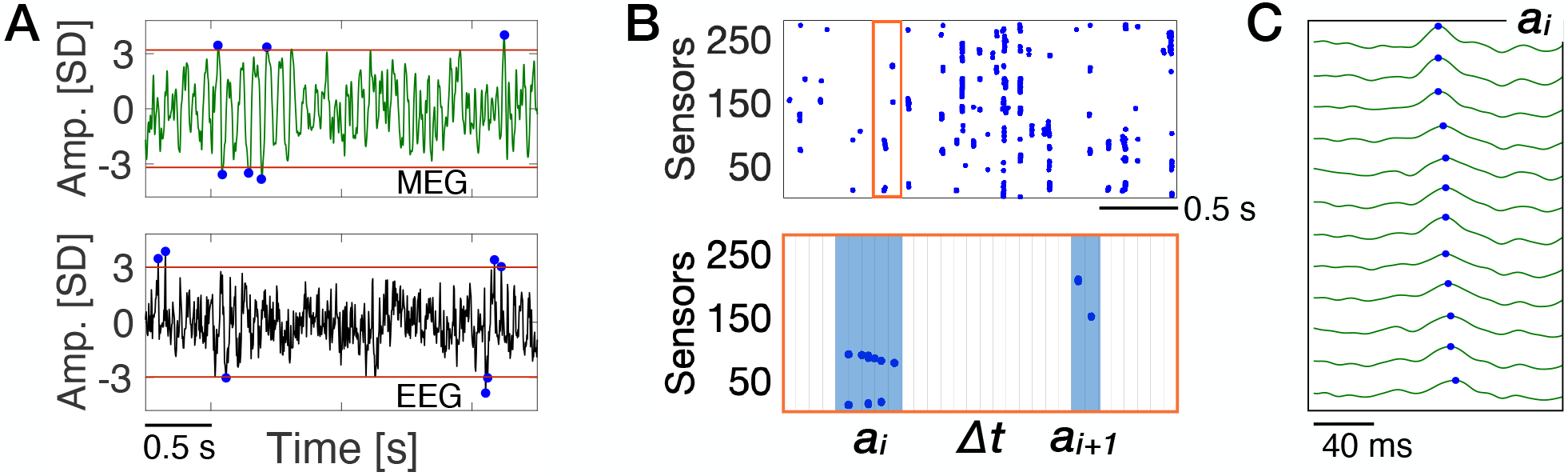
Neural activity cascades in the awake resting state. **(A)** Representative single-site MEG and EEG timeseries of resting-state activity in the human brain (top and bottom, respectively). Signals are z-normalized and the amplitude (Amp.) is in unit of standard deviation (SD) (STAR Methods). The most extreme point in each excursion beyond a threshold *h* (red lines) is treated as a discrete event (blue dots). **(B)** Representative raster of discrete events across all MEG sensors (n = 273, top). An avalanche *a*_*i*_ is defined as a sequence of temporal windows *ϵ* [*ϵ* = 2*T*_*meg*_ = 3.3 ms for the MEG (STAR Methods)] with at least one event in any of the sensors, preceded and followed by at least one window with no events in any of the sensors (bottom). The same procedure is used for the EEG (*ϵ* = *T*_*eeg*_ = 4 ms). **(C)** Temporal sequence of events (blue dots) that belong to the avalanche *a*_*i*_ in (B), which spreads over twelve different sensors. Events are presented in time ascending order, from top to bottom).

In Fig. 1B, we show a representative raster of events extracted from a MEG sensor array. The raster plot indicates that events tend to cluster in time across sensors—or cortical locations. We define a neural activity cascade, or avalanche, *a*_*i*_ as a sequence of consecutive time bins with at least one event in any of the sensors, preceded and followed by at least one empty time bin (Fig. 1B bottom, and Fig. 1C) [33]. To each avalanche, we associate a size, *s*_*i*_, which is given by the number of events occurring in the time bins that belong to it. It has been recently shown that the distribution *P*(*s*) of avalanche sizes follows a power-law behavior—*P*(*s*) ∝ *s*^−*τ*^—both in MEG and EEG recordings [36, 37, 17]. Power-law distributions indicate an absence of characteristic scales, a property in stark contrast with the main essence of brain rhythms, namely, characteristic times and amplitudes [2, 1, 3].

The scale-free power-law distribution of avalanche sizes does not suggest any potential link with neural oscillations. To characterize the dynamics of avalanches and identify relationships with coexisting rhythmic patterns, we turn our attention to the correlation properties of the cascading process. First, we analyze the autocorrelation function *C*(*t*) of the quantity *n*(*t*), the number of events occurring per unit time during the cascading process. We note that *n*(*t*) = 0 during quiet times, Δ*t*, which corresponds to periods with no threshold crossing events across the sensor array (Fig. 1B, bottom). We find that *C*(*t*) exhibits two distinct power-law regimes for *t <* 1 s, both in MEG and EEG recordings (Fig. 2A-B). At short time scales [regime (*A*_*<*_)], *C*(*t*) decays as 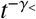; with *γ*_*<*_ ≃ 0.8; for 120 *< t <* 1000 ms [regime (*A*_*>*_)], *C*(*t*) shows a slower power law decay with *γ*_*>*_ ≃ 0.3 (Fig. 2A-B). Importantly, the crossover from regime (*A*_*<*_) to regime (*A*_*>*_) occurs around *t* ≃ 100 ms, which corresponds to the characteristic time of the alpha rhythm. The transition is preceded by a short plateau region between 20 – 100 ms (Fig. 2A-B). We observe that *C*(*t*) exhibits a further transition to a slower decaying regime around *t* = 1 s, which may be related to delta oscillations.

**Figure 2.**
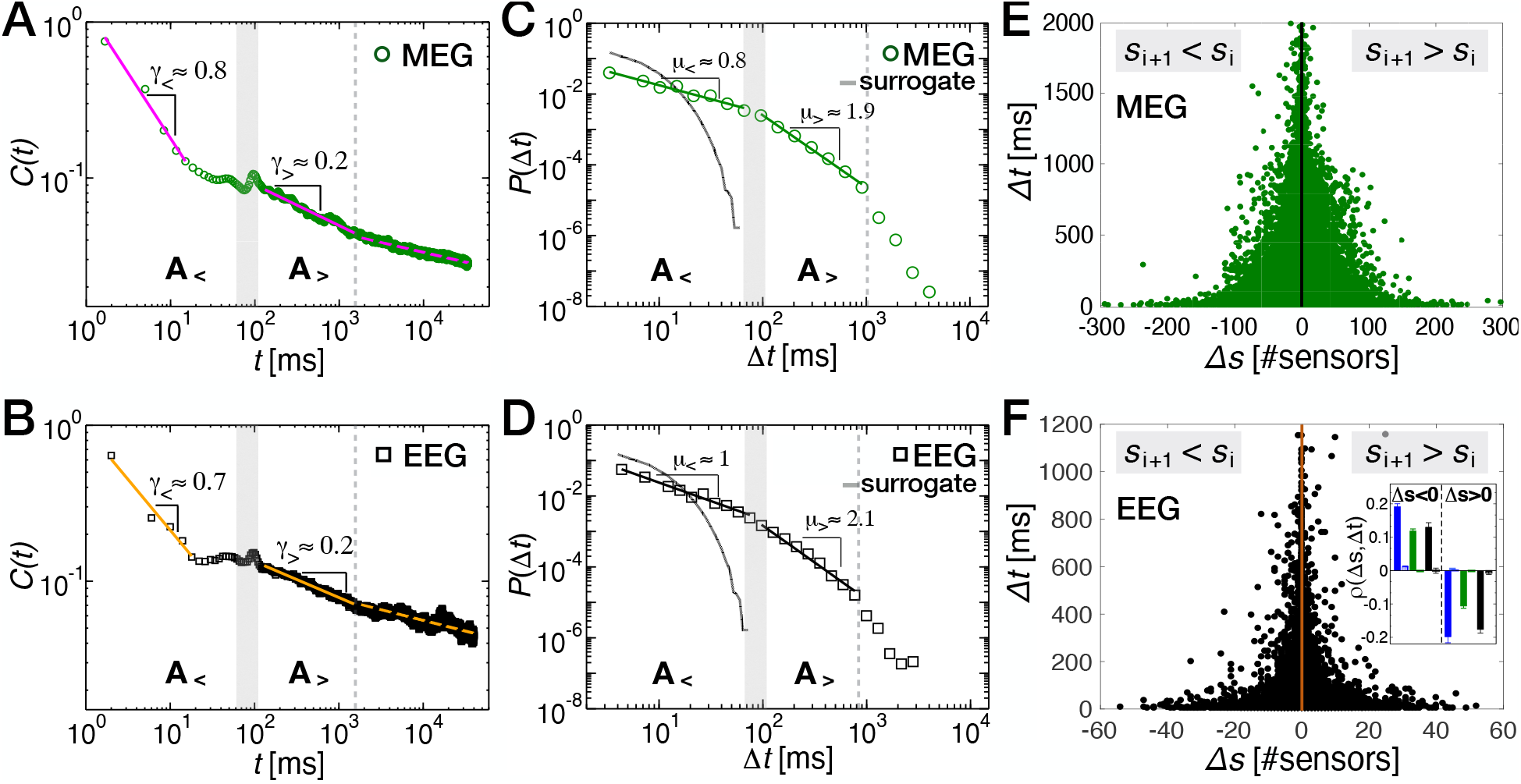
Transition in the dynamics of neural activity cascades at the time scale of the alpha rhythm during resting wake. **(A)** Auto-correlation *C*(*t*) of the network activity, *n*(*t*), in 4-min MEG recordings. *n*(*t*) is measured as the number of active sensors in each time bin *ϵ*. In the short time scales (regime (*A*_*<*_)), *C*(*t*) decays as a power-law with an exponent *γ*_*<*_ = 0.7633 0.0540. After a transition region (20 ms–120 ms) where *C*(*t*) is nearly constant and shows a small peak at *t* = 100 ms, the auto-correlation follows a power-law with an exponent *γ*_*>*_ = 0.2619 ± 0.0028 (regime (*A*_*>*_)). **(B)** Auto-correlation *C*(*t*) of the network activity for EEG recordings. Within regime (*A*_*<*_), *C*(*t*) decays as a power-law with an exponent *γ*_*<*_ = 0.6627 ± 0.0990, while within regime (*A*_*>*_) the auto-correlation follows a power-law with an exponent *γ*_*>*_ = 0.2309 ± 0.0019. **(C)** The distribution of quiet times *P*(Δ*t*) in 4-min MEG recordings (n = 70) shows a double power-law behavior with a regime (*A*_*<*_) for short Δ*t*’s characterized by an exponent *μ*_*<*_ = 0.7897 ± 0.0144, and a regime (*A*_*>*_) for longer Δ*t*’s with an exponent *μ*_*>*_ = 1.9690 ± 0.0394, followed by an exponential cutoff. The transition region (shaded area) between regime (*A*_*<*_) and (*A*_*>*_) is located around 100 ms. *P*(Δ*t*) from surrogate data (STAR Methods) shows an exponential behavior (grey curve). A similar behavior is observed in 40-min long MEG recordings (Fig. S3). **(D)** The distribution of quiet times in EEG recordings (n = 6) of the awake resting-state is consistent with the scenario described in (C): regime (*A*_*<*_), *μ*_*<*_ = 1.0832 ± 0.0544; regime (*A*_*>*_), *μ*_*>*_ = 2.0830 ± 0.1514. As in the MEG, the transition region (shaded area) is located at around 100 ms. *P*(Δ*t*) from surrogate data shows an exponential behavior (grey curve). **(E)** Scatter plot between Δ*s* and Δ*t* for all MEG subjects. Negative Δ*s*’s are positively correlated with their corresponding quiet times, whereas positive Δ*s*’s are anti-correlated with their relative quiet times (Spearman’s correlation coefficient: *ρ*(Δ*s <* 0, Δ*t*) = 0.1916 ± 0.0088 and *ρ*(Δ*s <* 0, Δ*t*) = 0.1188 ± 0.0061 in 40-min and 4-min MEG recordings, respectively; *ρ*(Δ*s >* 0, Δ*t*) = –0.1064 ± 0.0065 in 4-min MEG, *ρ*(Δ*s >* 0, Δ*t*) = –0.1985 ± 0.0181 in 40-min MEG). **(F)** The scatter plot between Δ*s* and Δ*t* for all EEG subjects exhibits the same behavior observed in the MEG (*ρ*(Δ*s <* 0, Δ*t*) = 0.1301 ± 0.0135; *ρ*(Δ*s >* 0, Δ*t*) = –0.1765 ± 0.0111). Inset: *ρ*(Δ*s*, Δ*t*) calculated separately for Δ*s <* 0 and Δ*s >* 0 (Blue: MEG 40 min; Green: MEG 4 min; Black: EEG). *ρ*(Δ*s*, Δ*t*) from surrogate data are plotted next to each bar, and are very close to zero in all cases. *t*-tests comparing original and surrogate data show that correlations between Δ*s* and Δ*t* are significant (*p*-value *<* 0.001; STAR Methods). Distributions were calculated using *ϵ* = 2*T*_*meg*_ = 3.3 ms and *ϵ* = 1*T*_*eeg*_ = 4 ms for MEG and EEG data, respectively. Results are independent of *ϵ* (Fig. S2). Power-law fits were performed using a maximum likelihood estimator, and compared with exponential fits via log-likelihood ratios (STAR Methods) [44, 45] (MEG 4-min; regime *A*_*<*_: *R* = 395, *p* = 0.03; regime *A*_*>*_, *R* = 346, *p* = 2 · 10^−18^. EEG; regime *A*_*<*_: *R* = 567, *p* = 4 · 10^−33^; regime *A*_*>*_: *R* = 72, *p* = 0.0003). In all cases, the power-law is more likely to describe the empirical data (STAR Methods). The *p*–value measures the significance of *R* and is defined in the STAR Methods.

The transition between distinct scaling regimes indicates a change in the cascading dynamics at the temporal scale of the alpha rhythm [46, 47]. We further investigate this point by analyzing the distribution of quiet times, *P*(Δ*t*) (Fig. 2C-D). Indeed, power-law decays in the auto-correlation *C*(*t*) can be related to power-laws in the quiet time distribution [48, 46, 47]. In both MEG and EEG data, we observe a close correspondence between short- and long-timescale regimes in *C*(*t*) and *P*(Δ*t*), with the transition from one another occurring consistently around 100 ms. Such a correspondence, as well as the hallmark of alpha oscillations on both *P*(Δ*t*) and *C*(*t*), is particularly evident in the analysis of individual subjects (Fig. S1). Specifically, for Δ*t <* 1 s, *P*(Δ*t*) is well described by two distinct power-law regimes with exponents *μ*_*<*_ and *μ*_*>*_, which correspond to (*A*_*<*_) and (*A*_*>*_) defined in Fig. 2A-B for the auto-correlation *C*(*t*). The crossover from one regime to the other is located around Δ*t* ≃ 100 ms, as in *C*(*t*) (Figs. 2C-D; see also Fig. S1-S3 for individual subjects and 40-min MEG). Thus, the regime (*A*_*<*_) includes the quiet times that are shorter than a single alpha cycle (about 100 ms), while (*A*_*>*_) includes the quiet times that span more than a single alpha cycle. This suggests that the change in the cascading dynamics marked by the crossover from (*A*_*<*_) to (*A*_*<*_) is closely connected to underlying properties of the alpha rhythm, as we shall investigate in the following sections. Such power-law regimes, (*A*_*<*_) and (*A*_*>*_), are followed by a faster decay of the probability *P*(Δ*t*) for Δ*t*’s longer than 1 s.

Next, we analyze the conditional distributions *P*(Δ*t* |*> s*_*c*_), where *s*_*c*_ is a minimum threshold on avalanche sizes. In short, one only considers the Δ*t*’s between consecutive avalanches of size *s > s*_*c*_. We examine *P*(Δ*t* |*s > s*_*c*_) for several *s*_*c*_ values. The analysis shows that, for Δ*t* = 100 ms, *P*(Δ*t*) is independent of *s*_*c*_, while *P*(Δ*t*) depends on *s*_*c*_ for both Δ*t <* 100 ms and Δ*t >* 100 ms (Figs. S4-S5). Specifically, for increasing *s*_*c*_ values, we find that (i) *P*(Δ*t*|*s > s*_*c*_) decreases for Δ*t <* 100 ms and the exponent *μ*_*<*_ decreases and tends to zero; (ii) *P*(Δ*t* |*s > s*_*c*_) increases for Δ*t >* 100 ms, and shows a power-law with decreasing exponents (Fig. S4). This shows that, in stark contrast to similar analyses in rats and zebrafish in absence of alpha oscillations [35, 49], Δ*t* = 100 ms behaves as a fixed point for the transformation that selectively removes avalanches smaller than a threshold *s*_*c*_, while the rest of the distribution changes. This further indicates that alpha oscillations play a key role in shaping the distribution of quiet times, and thus determining the emergent dynamics of avalanches in the resting state.

The double power-law, non-exponential quiet time distribution, as well as the autocorrelation *C*(*t*), implies that neural cascades of activity are strongly correlated [50, 51, 37, 52, 53, 54], and indicates that the nature of correlations in the cascading process depends on the time scales. Indeed, the distribution *P*(Δ*t*) calculated after random phase shuffling (STAR Methods) of the original brain signals is exponential (Figs. 2C, D). Furthermore, we observe that, despite the analogies between avalanches and earthquakes [55, 54], the double power-law behavior in *P*(Δ*t*) does not arise from the non-homogeneity of the avalanche rate (see Fig. S6), as found for earthquakes instead [56].

The observed functional behavior of *P*(Δ*t*), as well as the autocorrelation function *C*(*t*), suggests that the relationship between consecutive avalanches (and thus the underlying collective neural dynamics) undergoes a transition around Δ*t* ≃ 100 ms. Here, we hypothesize that this transition reflects a property of the neural dynamics associated with the generation and propagation of the alpha rhythm in resting-state brain activity. To verify this hypothesis, we investigate whether the collective neural dynamics encoded in avalanche characteristics (e.g. their sizes) exhibits signatures of dynamic transitions that correlate with alpha oscillations.

### Attenuation-amplification dynamics (AAD) of neural activity in the awake resting-state

To this end, we consider the avalanche size increments Δ*s* = *s*_*i*+1_ – *s*_*i*_ between the size of avalanche *a*_*i*_ and the size of the subsequent avalanche *a*_*i*+1_, and analyze their relationship with the corresponding quiet times Δ*t*_*i*_ (Fig. 1). Negative values of Δ*s* imply attenuation, i.e. the avalanche *a*_*i*+1_ is smaller than its preceding one *a*_*i*_, whereas positive values imply amplification, namely, the avalanche *a*_*i*+1_ is larger than the preceding avalanche *a*_*i*_

We first examine the scatter plot between Δ*s*’s and the corresponding Δ*t*’s (Figs. 2E,F). For Δ*s <* 0, we observe that large negative Δ*s*’s occur with short quiet times Δ*t*’s and vice versa, both in MEG (Fig. 2E) and EEG recordings (Fig. 2F). The corresponding value of the Spearman’s correlation coefficient *ρ*(Δ*s <* 0, Δ*t*) (STAR Methods) is significantly larger than zero (Fig. 2F, inset), indicating that negative size increments are positively correlated with their corresponding quiet times, i.e. the more negative the increment, the shorter the quiet time separating two avalanches. In contrast, we find that positive size increments, Δ*s >* 0, tend to be significantly anti-correlated with the corresponding quiet times both in MEG and EEG data (Fig. 2F, inset).

To dissect the dependency of size increments on the time lag separating consecutive avalanches and identify the connection with the alpha rhythm, we scrutinize the correlation landscape hidden in the density distribution across the Δ*t*Δ*s* plane of the scatter plots shown in Figs. 2E and F. To this aim, we systematically compare the spatial structure of the density in the plane Δ*t*Δ*s* with surrogate densities obtained by randomly reshuffling avalanche sizes. In this way, both the distribution and the temporal order of quiet times are preserved, as well as the size distribution (Fig. 3A).

**Figure 3.**
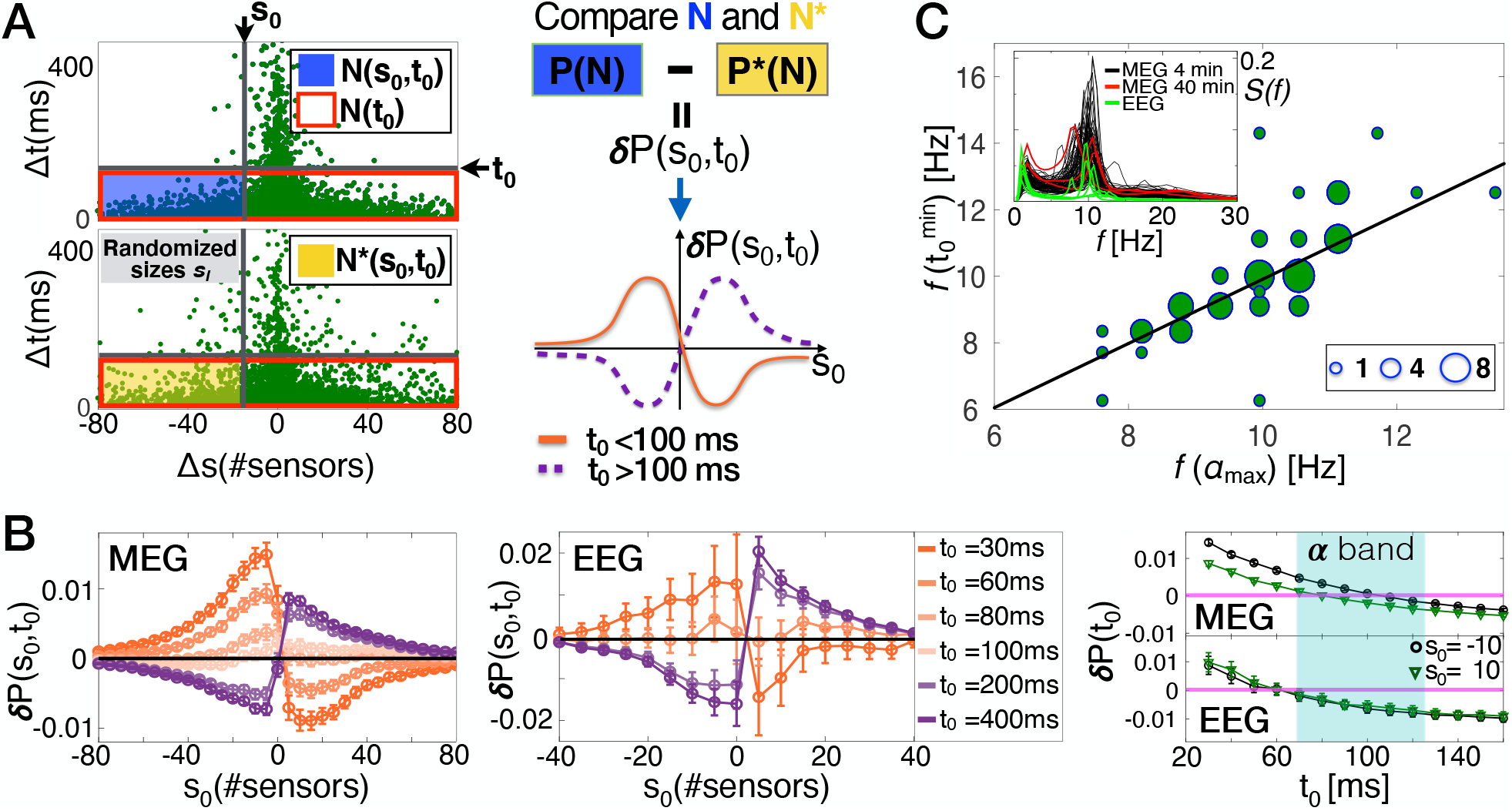
The monotonic relationship between consecutive avalanches undergoes a transition from an attenuation to an amplification regime that correlates with the characteristic time of the alpha rhythm. **(A)** Schematic definition of the quantity *δP*(*s*_0_, *t*_0_) ≡ *δP*(Δ*s < s*_0_|Δ*t < t*_0_). **(Left panel)** The conditional probabilities *P*(*s*_0_, *t*_0_) and the surrogate conditional probabilities *P*^∗^(*s*_0_, *t*_0_) are proportional to the number of points *N*(*s*_0_, *t*_0_) and *N*^∗^(*s*_0_, *t*_0_) in the region *R*(*s*_0_, *t*_0_) of the plane Δ*s*Δ*t* defined by the thresholds *s*_0_ and *t*_0_ (blue and yellow rectangles, respectively). **(Right panel, top)** The quantity *δP*(*s*_0_, *t*_0_) compares the original and the surrogate density in the region *R*(*s*_0_, *t*_0_) of the plane, and is defined as the difference between *P*(*s*_0_, *t*_0_) and *P*^∗^(*s*_0_, *t*_0_), the conditional probabilities associated with the density *N* and *N*^∗^ for original and surrogate data (STAR Methods). (**Right panel, bottom**) The quantity *δP*(*s*_0_, *t*_0_) as a function of *s*_0_ for a given threshold *t*_0_ on Δ*t*’s exhibits two relevant scenarios: For *t*_0_ *<* 100 ms (attenuation regime, orange thick line), *δP*(*s*_0_, *t*_0_) has a maximum for *s*_0_ *<* 0, implying that *s*_*i*+1_ tends to be smaller than *s*_*i*_ for Δ*t < t*_0_; for *t*_0_ *>* 100 ms (amplification regime, purple dashed line), *δP*(*s*_0_, *t*_0_) has a maximum for *s*_0_ *>* 0, implying that, for Δ*t < t*_0_, *s*_*i*+1_ tends to be larger than *s*_*i*_. **(B)** *δP*(*s*_0_, *t*_0_) as a function of the threshold *s*_0_ on Δ*s* for different values of the threshold *t*_0_ on Δ*t*’s for 4-min MEG (left panel, n = 70) and EEG (middle panel, n = 6) data. The error bar on each data point is two times the standard deviation *σ*^∗^ associated with the surrogates *P*^∗^(*s*_0_, *t*_0_) (see STAR Methods). For a given *t*_0_, the maximum in *δP*(*s*_0_, *t*_0_) indicates the preferred relation between consecutive avalanches separated by quiet times shorter than *t*_0_. Both in MEG and EEG recordings (left and middle panel), for *t*_0_ smaller than ≈100 ms, *δP*(*s*_0_, *t*_0_) has a maximum at *s*_0_ *<* 0, and thus an avalanche tends to be smaller than its preceding one (*s*_*i*+1_ *< s*_*i*_, attenuation regime). On the contrary, for *t*_0_ larger than 100 ms, the maximum moves towards positive *s*_0_, implying that a given avalanche tends to be larger than the preceding one (*s*_*i*+1_ *> s*_*i*_, amplification regime). For *t*_0_ ≈ 100 ms, *δP*(*s*_0_, *t*_0_) is very close to zero for each *s*_0_, indicating that at *t*_0_ ≃ 100 ms, there is not a preferred sign for Δ*s*, and that Δ*t* = 100 ms is a transition point from one dynamical regime to another—an attenuation-amplification transition. A similar behavior is observed in 40-min MEG recordings (Fig. S7). (Right panel) *δP*(*s*_0_, *t*_0_) as a function of *t*_0_ for *s*_0_ = –10 and *s*_0_ = 10 in 4-min MEG and EEG. For each *s*_0_, *δP*(*t*_0_) transitions from positive to negative values in a range of *t*_0_ corresponding to the alpha band (≈80 ms and≈130 ms). A similar behavior is observed in 40 min MEG (Fig. S7). **(C)** The scatter plot between *f* (*α*_*max*_) and 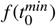 shows that the attenuation-amplification transition in *δP*(*s*_0_, *t*_0_) correlates with the maximum in the *α* band of the power spectrum. The size of each circle is proportional to the number of points (subjects) in the corresponding region of the plane. The black thick line is a linear fit, *Y* = *A* · *X* + *B*, with *A* = 0.96 and *B* = 0.26 (*r*^2^ = 0.80). Inset: Power spectrum *S* (*f*) for each subject (black lines, MEG 4 min; red lines, MEG 40 min; green lines, EEG).

The local density distribution resulting from the avalanche process can be described using the conditional probability *P*(*s*_0_, *t*_0_) ≡ *P*(Δ*s < s*_0_|Δ*t < t*_0_) = *N*(Δ*s < s*_0_|Δ*t < t*_0_)*/N*(Δ*t < t*_0_), where the thresholds *s*_0_ and *t*_0_ delimit the plane region *R*(*s*_0_, *t*_0_) under investigation (Fig. 3A, top left, blue shaded area). An analogous quantity *P*^∗^(*s*_0_, *t*_0_) can be associated with the surrogate region resulting from the size reshuffling procedure (Fig. 3A, bottom left, yellow shaded area). The difference *δP*(*s*_0_, *t*_0_) ≡ *δP*(Δ*s < s*_0_|Δ*t < t*_0_) = *P*(*s*_0_, *t*_0_) –*P*^∗^(*s*_0_, *t*_0_) is a measure of the likelihood that a specific distribution of points in the region *R*(*s*_0_, *t*_0_) results from the actual avalanche dynamics, rather than from random avalanche occurrences, and is used to examine the relationship between sizes of consecutive avalanches as a function of the quiet times separating them. We indicate with *σ*^∗^ the standard deviation associated with the surrogates *P*^∗^(*s*_0_, *t*_0_). Thus, if|*δP*(*s*_0_, *t*_0_)| *>* 2*σ*^∗^, *P*(*s*_0_, *t*_0_) and *P*^∗^(*s*_0_, *t*_0_) are significantly different (*p*-value *<* 0.05; see STAR Methods), and the distribution of points in the region *R*(*s*_0_, *t*_0_) is considered to reflect a specific, non-random relationship between Δ*s* and Δ*t*.

We study *δP*(*s*_0_, *t*_0_) as a function of *s*_0_ for fixed *t*_0_ values. For each fixed value of *t*_0_, there are two possible relevant scenarios for the function *δP*(*s*_0_, *t*_0_) (Fig. 3A, right): (i) *Attenuation regime. δP*(*s*_0_, *t*_0_) is positive and monotonically increases for –∞ *< s*_0_ *<* 0, reaches a maximum, and then decreases, taking a minimum at some *s*_0_ *>* 0 (Fig. 3A, right, bottom, orange thick line). In this regime, the size increments, Δ*s* = *s*_*i*+1_ – *s*_*i*_, tend to be negative, and the avalanche *a*_*i*+1_ tends to be smaller than the preceding avalanche *a*_*i*_; (ii) *Amplification regime. δP*(*s*_0_, *t*_0_) is negative and monotonically decreases for –∞ *< s*_0_ *<* 0, reaches a minimum, and then increases, taking a local maximum at some *s*_0_ *>* 0 (Fig. 3A, right, bottom, purple dashed line). In the latter regime, the amplification regime, the size increments tend to be positive, and a given avalanche *a*_*i*+1_ tends to be larger than its preceding avalanche *a*_*i*_. Following this approach, we show that the attenuation and amplification regimes correspond to regimes (*A*_*<*_) and (*A*_*>*_) defined in Fig. 2.

We first analyze *δP*(*s*_0_, *t*_0_) as a function of *s*_0_ for several fixed values of the threshold *t*_0_ (Fig. 3B). If the relation between consecutive avalanches depends on their distance in time, then *δP*(*s*_0_; *t*_0_) changes as we consider different *t*_0_ values. In Fig. 3B we show the function *δP*(*s*_0_, *t*_0_) evaluated for *t*_0_ values ranging between 30 ms and 1000 ms. We find that *δP*(*s*_0_; *t*_0_) follows the attenuation regime for *t*_0_’s smaller than approximately 100 ms, whereas it conforms to the amplification regime for larger *t*_0_ values. Thus, size increments Δ*s* tend to be negative if the quiet time between consecutive avalanches is shorter than 100 ms, and positive otherwise. This behavior is also observed in 40-min MEG data (Fig. S7), and is consistent across subjects (Fig. S8). The tendency for negative Δ*s*’s to be coupled with Δ*t <* 100 ms implies that avalanche sizes preferentially exhibit a decreasing trend on short time scales, a clear sign of attenuated bursting activity. On the other hand, the significant likelihood for positive increments at longer time scales suggests the presence of a regulatory mechanism that, after discharging cycles with decreasing avalanche sizes, amplifies bursting activity and leads to the appearance of larger avalanches. The transition from the attenuation to the amplification regime occurs at *t*_0_ ≈ 100 ms, where we observe that *δP*(*s*_0_, *t*_0_) ≃ 0 (Fig. 3B). Importantly, *δP*(*s*_0_, *t*_0_) ≃ 0 implies that *P*(*s*_0_, *t*_0_) ≃ *P*^∗^(*s*_0_, *t*_0_), which in turn implies that, in the plane Δ*t*Δ*s*, the local density around Δ*t* ≈ 100 ms is comparable to the density obtained with the reshuffled avalanche sizes. Crucially, we observe that the attenuation and amplification regimes correspond to regime (*A*_*<*_) and (*A*_*>*_) in the quiet time distributions, and the attenuation-amplification transition coincides with the crossover from regime (*A*_*<*_) to regime (*A*_*>*_) (Fig. 2C-D).

### The AAD of the awake resting-state correlates with the alpha rhythm

To identify the transition point *t*^*min*^ from the attenuation to the amplification regime, we study *δP*(Δ*s < s*_0_, Δ*t < t*_0_) as a function of *t*_0_ for a range of fixed *s*_0_ values — i.e. *δP*(*t*_0_) (Fig. 3B, right). We plot *δP*(*t*_0_) for *s*_0_ = ±10, where the quantity *δP*(Δ*s < s*_0_, Δ*t < t*_0_) is generally non-zero away from the transition point (Fig. 3, left and middle panels). A similar behavior is observed for other *s*_0_ values (Fig. S7). We find that the transition from positive to negative *δP*(*t*_0_) lies mostly in the range [70 ms, 130 ms] (Fig. 3B and S7), which approximately corresponds to oscillations in the frequency range [8–13] Hz, commonly identified as the alpha band. Similar results are obtained in 40-min MEG recordings (Fig. S7). Then we define 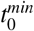 as the *t*_0_ that minimizes |*δP*(Δ*s < s*_0_|Δ*t < t*_0_)| over a range of relevant *s*_0_ values, namely 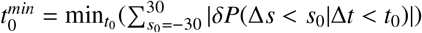.

Next, we show that the attenuation-amplification transition correlates with the timescale characteristic of the alpha rhythm, and is consistent across subjects in both MEG and EEG recordings of the awake resting-state (Fig. S7-S8). In Fig. 3C, for all subjects, we plot the frequency *f* (*α*_*max*_), which corresponds to the maximum power in the alpha band, versus the transition frequency 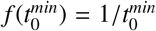 corresponding to the transition point 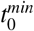 from the attenuation regime to the amplification regime (Fig. 3C). We observe that the frequency 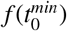 correlates with *f* (*α*_*max*_) (*r*^2^ = 0.80), indicating that the attenuation-amplification transition is intimately connected to the dynamics of alpha oscillations. Importantly, the correlation between the transition frequency 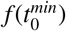 and the *f* (*α*_*max*_) suggests that the crossover both in the distribution of quiet times and in the quantity *δP*(*s*_0_, *t*_0_) relates to a basic difference in the cascading process within and between alpha cycles.

### The alpha “waxing and waning”: a long-term mechanism that regulates attenuation and amplification bouts according to the Omori law

We have shown that the AAD of the resting-state cascading process acts on a timescale of less than a second, with an attenuation-amplification transition that correlates with the characteristic frequency of alpha oscillations. Importantly, alpha waves are also known to exhibit long-term alternation between higher and lower amplitude fluctuations over the timescales of several seconds—the so called “waxing and waning” phenomenon [19, 20, 18]. To characterize the resting-state cascading process in relation to alpha “waxing and waning”, we focus on large cascades of activity, which we define as avalanches larger than a given size *s*^∗^ and label them as main avalanches, *A*^∗^ (Fig. 4A, left top panel). Large avalanches are synchronous events consisting of time-clustered, higher amplitude signal fluctuations over a large number of sensors—see the raster plot in Fig. 4A, left bottom panel. We observe that, following a main avalanche *A*^∗^ (identified by blue arrows in the right panel of Fig. 4A), the sizes of activity cascades tend to rapidly decrease, with fluctuations that also decrease with the time elapsed from *A*^∗^. We refer to the activity following a main avalanche as the Omori sequence (Fig. 4A, left panel), in analogy with the sequence of aftershocks that follows a main earthquake.

**Figure 4.**
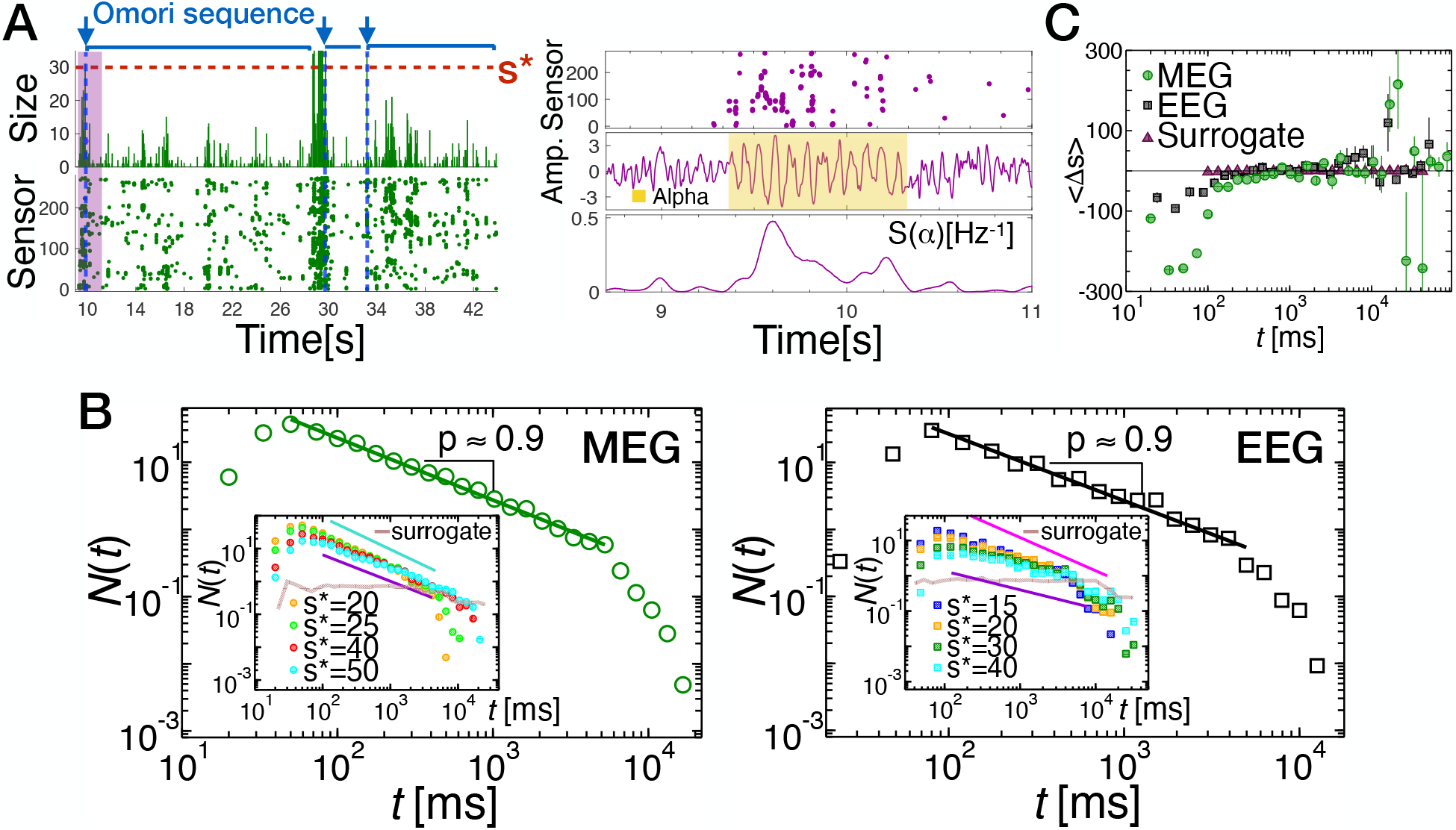
The alpha “waxing and waning” is described by the Omori law in the underlying neural cascading process. **(A)** A raster plot of 35 s of activity (left, bottom) and the corresponding avalanche sizes (left, top) showing three Omori sequences of different durations ranging between about 2 s and 15 s and including several alpha cycles. Each sequence starts with a main avalanche *A*^∗^ with a size larger than 30 (blue arrow). Activity decreases over time until the next main avalanche. Raster plot of the start of the first Omori sequence highlighted in magenta in the left panel (right, top). Brain activity around the main avalanche corresponds to high-amplitude alpha bursts (right, middle panel; yellow shaded area), preceded and followed by lower-amplitude fluctuations. The instantaneous power at 10 Hz peaks around the main avalanche, and then rapidly decreases (right, bottom). **(B)** *N*(*t*) as a function of the time *t* elapsed after a main avalanche *A*^∗^ in 4-min MEG resting brain activity (left, individual subject). *N*(*t*)decreases as the reciprocal of the time elapsed from the main avalanche *A*^∗^, i.e. *N*(*t*) ∝ *t*^−*p*^ with *p* ≃ 1 (*p* = 0.9240 ± 0.0169. *s*^∗^ = 30; *s*∗ */*(#*MEGsensors*) ≃ 0.11). Inset: *N*(*t*) for different values of the threshold *s*^∗^ defining a main avalanche *A*^∗^. The exponent *p* describing the power-law decay of *N*(*t*) ranges between 0.9773 ± 0.0302 (turquoise line) and 0.8054 ± 0.020 (violet line). Similar results are obtained in 40-min MEG recordings (Fig. S9). The Omori-law is verified in the EEG of the resting state (right, individual subject), with *p* = 0.9384 ± 0.0321 for *s*^∗^ = 10 (*s*^∗^*/*(#*EEGsensors*) ≃ 0.16). Power-law fits were performed using a maximum likelihood estimator, and compared with exponential fits via log-likelihood ratios (STAR Methods) [44] (MEG 4-min: *R* = 270, *p*-value = 4 · 10^−17^. EEG: *R* = 70, *p*-value = 3 · 10^−5^). The p-*value* measures the significance of *R* and is defined in the STAR Methods. In all cases, the power-law is more likely to describe the empirical data. Inset: *N*(*t*) for different values of the threshold *s*^∗^ defining a main avalanche *A*^∗^. The exponent *p* describing the power-law decay of *N*(*t*) ranges between 0.9384 ± 0.0321 (magenta line) and 0.5289 ± 0.0471 (violet line). *N*(*t*) is independent of *t* in the surrogate data (insets, brown curves). **(C)** Average Δ*s* = *s*_*i*+1_ – *s*_*i*_ between consecutive avalanche occurring within two main avalanches as a function of the time *t* elapsed from the first main avalanche (4-min MEG: n = 70; EEG: n = 6). Error bars represent the standard error of the mean. We note that for surrogate data, Δ*s* is always close to zero, independently of *t* (brown triangles).

Such dynamics follow the attenuation-amplification principle over timescales ranging between a few seconds to a few tens of seconds, and is strongly reminiscent of the alpha “waxing and waning” [20]. Following this analogy, we analyze the brain activity around the main avalanches *A*^∗^’s. In Fig. 4A (right), we show the analysis of a raster plot segment from an Omori sequence (marked in magenta in the left panel). We observe that the main avalanche identified in the raster plot (top) corresponds to higher amplitude alpha bursts (middle), with large peaks in the alpha power *S* (*α*) (bottom). Furthermore, we notice that before and after the main avalanche, both the signal amplitude and the alpha power decrease considerably (Fig. 4A, middle and bottom).

Next, we proceed to quantify the Omori sequences by analyzing the number *N*(*t*) of avalanches per unit time occurring after a time *t* has elapsed from the main avalanches. We find that *N*(*t*) decays as *t*^−*p*^ across a wide range of *t*’s in both 4-min MEG and EEG recordings (Fig. 4B). We observe that the exponent *p* is close to one for a range of threshold values *s*^∗^ used to identify the main avalanches, and its value decreases when the threshold *s*^∗^ becomes too large (Fig. 4B, insets). This behavior is also observed in 40-min MEG recordings, and is consistent across subjects (Fig. S9). The power-law decay *N*(*t*) ∝ *t*^−*p*^ of the number of events following a main avalanche is consistent with the generalized Omori law, *N*(*t*) ∝ (*t* + *c*)^−*p*^ (where *c* is a parameter related to the onset of the power-law regime), which describes the temporal organization of aftershocks following a main earthquake [50, 51]. The presence of the Omori law indicates that, after a main avalanche, the occurrence of the following avalanches is correlated over a wide range of timescales—up to several seconds, the location of the power-law cutoff. In particular, the power-law decay, *N*(*t*) ∝ *t*^−1^, implies that the activity after a large cascade is characterized by temporal clustering over unusually long timescales—up to tens of seconds—and the alternation between high- and low-amplitude fluctuations does not have characteristic temporal scales [18]. In analogy with the earthquake dynamics, the Omori law for avalanches may be related to the slow build up and discharge of synaptic resource across neural populations following main avalanches. Importantly, we observe that, within an Omori sequence, the average Δ*s* between consecutive avalanches is always negative for *t <* 100 ms (Fig. 4C). For *t* longer than a few hundreds of milliseconds, the average ⟨Δ*s*⟩ tends to transitions to positive values, although larger statistical samples would be needed to make a robust assessment of Δ*s* in this region.

### Neural activity cascades during sleep do not obey attenuation-amplification dynamics

We have demonstrated that collective neural activity during resting wakefulness is characterized by AAD, namely, the alternation of two distinct dynamical regimes: an attenuation regime within the alpha cycle, and an amplification regime between alpha cycles (Fig. 3). The AAD is part of the Omori-type dynamics encompassing resting-state brain activity over timescales of several seconds. To verify that these characteristics of the cascading dynamics are specific to the awake resting-state dominated by the alpha rhythm, we next investigate the dynamics of neural cascades during NREM sleep. In contrast to the awake resting-state, brain activity during NREM sleep shows a limited amount of alpha oscillations and instead is largely dominated by slow oscillations in the delta band (1–4 Hz) [4].

We consider EEG recordings across approximately 8 h of night sleep (STAR Methods), and analyze the distribution of quiet times between consecutive avalanches. We observe that, unlike the awake resting-state, the distribution of quiet times during NREM sleep exhibits a single power-law regime with an exponent *μ*_*s*_ ≃ 1.8, followed by an exponential decay for Δ*t >* 10 s (Fig. 5A). Correspondingly, the auto-correlation function *C*(*t*) of the instantaneous activity, defined as the sum of the absolute values of all signals exceeding the threshold in a time bin, shows a power-law decay with an exponent *γ* ≃ 0.3 for 1 s *< t <* 10 s, followed by a slower decay for larger *t*’s (Fig. 5B). Unlike in the awake resting-state (Fig. 2), during NREM sleep, *C*(*t*) exhibits an exponential decay at shorter timescales (*t <* 500*ms*) (Fig. 5B).

**Figure 5.**
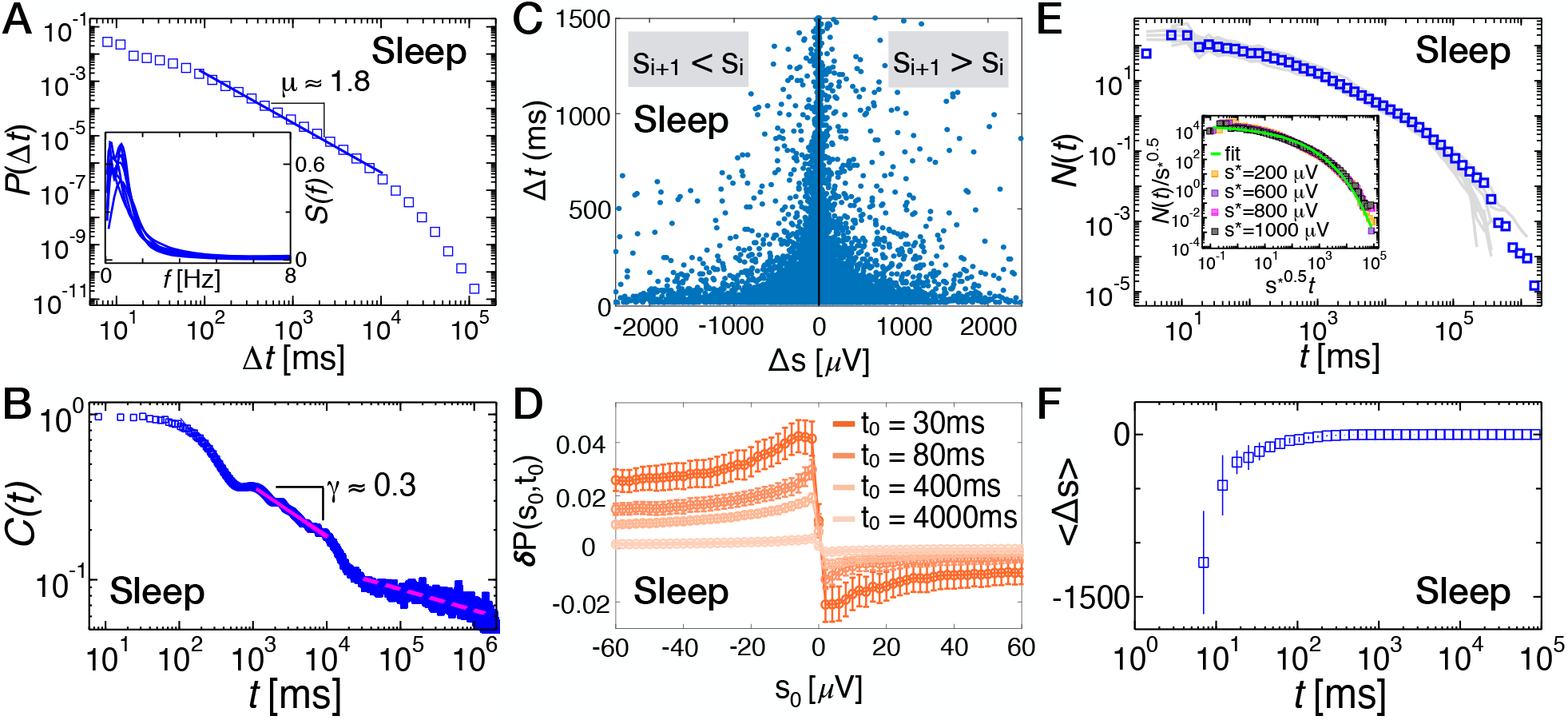
Neural activity cascades during sleep do not exhibit AAD and do not obey the Omori law. **(A)** Distributions of quiet times *P*(Δ*t*) between consecutive avalanches during sleep (pooled data; n = 10). Unlike the resting state, during sleep, quiet time distributions show a single power-law regime characterized by an exponent *μ* = 1.8311 ± 0.0018 (*R* = 107; STAR Methods) and followed by an exponential cutoff. Inset: Power spectra for individual subjects. The maximum is always in the range 0.5–2 Hz, within the delta band. **(B)** Auto-correlation *C*(*t*) of the instantaneous activity measured in each time bin *ϵ*. For *t <* 500 ms, *C*(*t*) exhibits an exponential decay, which is followed by a plateau between 500 ms and 1,000 ms, corresponding to 1–2 Hz delta oscillations. For 10^3^ *< t <* 10^4^ ms, *C*(*t*) decays as a power-law with an exponent *γ* = 0.3017 ± 0.0011. **(C)** Scatter plot between Δ*s* and Δ*t* during sleep (all subjects). Negative Δ*s*’s are positively correlated with their corresponding quiet times, whereas positive Δ*s*’s are anti-correlated with their relative quiet times. The Spearman’s correlation coefficient *ρ*(Δ*s*, Δ*t*) calculated separately for Δ*s <* 0 and Δ*s >* 0. *ρ*(Δ*s*, Δ*t*) is positive for Δ*s <* 0 and negative for Δ*s >* 0: *ρ*(Δ*s <* 0, Δ*t*) = 0.1627 ± 0.0032; *ρ*(Δ*s >* 0, Δ*t*) = –0.1022 ± 0.0041. **(D)** The quantity *δP*(*s*_0_, *t*_0_) as a function of *s*_0_ for different values of the threshold *t*_0_ on Δ*t*’s. The error bar on each data point is 2*σ*^∗^ (*σ*^∗^ is the SD associated with the surrogates *P*^∗^ (STAR Methods)). For each value of *t*_0_, *δP* is always positive for *s*_0_ *<* 0 and negative for *s*_0_ *>* 0, and takes its maximum (minimum) at *s*_0_ ≃–10 (*s*_0_ ≃10). Hence, for successive avalanches, the following avalanche tends to be smaller than the preceding one (attenuation regime), independently of the quiet time that separates them (cf. Fig. 3). **(E)** Number of avalanches per unit time, *N*(*t*), occurring after a main avalanche, i.e. an avalanche of size larger than *s*^∗^ (Grey lines: Individual subjects; Symbols: Pooled data). Unlike the resting wake, *N*(*t*) does not obey the Omori law, but is well fitted by a stretched exponential 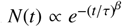. Inset: *N*(*t*) for different values of the threshold *s*^∗^ used to define a main avalanche. Data for different *s*^∗^’s collapse onto a single curve when *t* is rescaled by *s*^∗0.5^. Green line: Stretched exponential fit 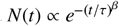, with *β* = 0.25. **(F)** The average Δ*s* = *s*_*i* 1_ – *s*_*i*_ between consecutive avalanches occurring within two main avalanches is always negative, and monotonically increases with *t*, the time elapsed from the main avalanche. Error bars represent the standard error of the mean.

Next, we examine the scatter plot between the Δ*s*’s and the corresponding Δ*t*’s (Fig. 5C). We find that large negative Δ*s*’s occur with short quiet times Δ*t*’s, while positive size increments tend to be anti-correlated with the corresponding quiet times (Fig. 5C). This behavior is similar to the behavior we observed during resting wakefulness (Figs. 2E, F). However, the relationship between consecutive avalanches as a function of the time separation exhibits a rather different behavior during NREM sleep. We analyze the quantity *δP*(*s*_0_; *t*_0_) defined in Fig. 3 as a function of *s*_0_ for a range of *t*_0_ values between 30 ms and 4,000 ms (Fig. 5D). We find that *δP*(*s*_0_; *t*_0_) follows the attenuation regime defined in Fig. 3 for all *t*_0_ values (Fig. 5D), namely, the size increments Δ*s* between consecutive avalanches tend to be always negative. This implies that avalanche sizes preferentially exhibit a decreasing trend, an attenuation effect that is particularly strong for Δ*t*’s shorter than 400 ms. Such a behavior is in stark contrast with our observations during resting wakefulness, where we found a transition from an attenuation regime—Δ*s <* 0 for Δ*t <* 100 ms—to an amplification regime—Δ*s >* 0 for Δ*t >* 100 ms—at the characteristic time of the alpha rhythm (Fig. 3).

We next analyze the number *N*(*t*) of avalanches per unit time occurring after a time *t* has elapsed from a main avalanche *A*^∗^. We find that, unlike in the awake resting-state, *N*(*t*) decays according to a stretched exponential, that is 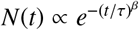 with *β* = 0.25. This behavior is consistent across subjects (Fig. 5E, grey curves), and does not depend on the threshold *s*^∗^ used to define the main avalanches *A*^∗^, as demonstrated by the data collapse in the inset of Fig. 5E. Furthermore, in contrast to the awake resting-state, we find that the average Δ*s* in the sequence of avalanches between two main shocks is always negative, and slowly approaches zero for *t >* (1 – 2 s) (Fig. 5F).

## Discussion

In this paper, we provided a description of resting-state brain activity that uncovers the dynamic organization of neural activity cascades in relation to brain rhythms, and offers new insights into the functional role of alpha oscillations in the awake resting-state. Our analysis shows that the collective neural dynamics underlying resting-state brain activity is characterized by the rhythmic alternation of attenuation-amplification bouts, which is modulated by the alpha rhythm across multiple timescales. On a timescale of a few alpha cycles (*<* 1 s), attenuation of neural activity cascades is found within the typical length of the alpha cycle, i.e. about 100 ms [2, 1], while amplification of neural activity cascades occurs over timescales of a few hundreds of milliseconds. Significantly, the attenuation-to-amplification transition consistently correlates with the dominant frequency in the alpha band. We have shown that these short-term AAD is part of a large-scale, size-dependent temporal structure of neural cascades that obeys the Omori law [50]: large avalanches (main avalanches) are followed by increasingly smaller avalanches at a rate that decays as a power-law of the time elapsed from the main avalanche—a long-term AAD regulating brain activity over the timescale of seconds. Importantly, both the short-term AAD and the Omori law are unique to the awake resting-state, and are not present during NREM sleep.

The dynamic structure of neural cascades during the awake resting-state contains the hallmarks of two key functional characteristics of the alpha rhythm: (i) The timing of inhibition and excitation in cortical networks, and (ii) The fluctuations in amplitude known as “waxing and waning” [19, 42, 20, 57, 6, 58, 18]. The short-term AAD reported in the present study indicates that inhibition may be activated to gradually suppress the cascading process within about 100 ms during the attenuation regime, while excitability is successively enhanced to amplify neural cascades over the timescales of a few alpha cycles (a few hundreds of milliseconds). Coherently, at the crossover between attenuation and amplification, i.e. at about the 100 ms characteristic of alpha waves, there is no clear monotonic relation between consecutive avalanches, which is consistent with a random organization of avalanche sizes—a clear transition signature. This is consistent with the hypothesis that alpha-mediated inhibition is applied in rhythmic cycles, a mechanism referred to as “pulsed inhibition” [30, 31, 8]. At the same time, our findings also indicate an active role for alpha waves in timing the increase in network excitability (increased excitation or dis-inhibition), an effect not previously described. This suggests a dual role for alpha oscillations, going beyond the pulsed inhibition hypothesis. Furthermore, our analysis shows that the attenuation-amplification principle governs resting-state brain activity across scales, revealing a precise structure in the cascading process underlying the long-term “waxing and waning” phenomenon [20]. We found that the high-amplitude alpha bursts function as main inhibitory events. Between such events—which we call main avalanches—smaller and increasingly sparser cascades occur, obeying the short-term AAD. These observations indicate that the precise short-term dual role—attenuating and amplifying—of the alpha rhythm is embedded in a size-time dependent long-term organization captured by the Omori law. The range of variability implied by the power-law behavior of the Omori law, previously known as “waxing and waning” phenomena with different timescales [20], indicates that the fluctuations associated to the “waxing and waning” of the alpha rhythm do not have a characteristic time, as recent studies also suggested [18, 26]. This may provide the brain with the flexibility necessary to organize complex streams of information, while maintaining precise information control through timely short-term AAD.

Such findings point to an intermittent rather than periodic nature of the alpha waxing and waning. This is in line with recent analyses of the resting-state brain activity [18, 26] showing that the power in the alpha band follows a bistable distribution, with large-scale high- and low-power modes. Single sensor has shown that the activity switches between high- and low-power modes [18]. Both the dwell time in high- and low-power modes are distributed according to a stretched exponential, indicating that the alternation between modes is bursty, or erratic, in nature, rather than periodic. We notice that, unlike neuronal avalanches—spatio-temporal events unfolding over multiple sensors across multiple time bins—, these quantities are defined on single sensor signals. Moreover, avalanches are a collective measure, and, as such, their dynamics arises from the complex relationship among all sensor signals (i.e., large and distinct populations of neurons). However, the building blocks of neuronal avalanches, i.e. excursions over threshold in individual signals, obey Weibull-like statistics [23], indicating that, when constrained to the individual sensors signals that participate in the avalanches, there are close analogies with the results in [18].

In stark contrast with our observations in the awake resting-state, we found that the cascading process during NREM sleep, where the alpha rhythm is nearly absent, does not show AAD, and the avalanche occurrence rate after a large avalanche follows a stretched exponential decay. The stretched exponential can be understood as a superposition of exponential decays with different characteristic times. In this case, the cascading process can be seen as the superposition of many independently acting entities, each with a specific fixed cascading rate. Thus, the presence of a stretched exponential decay during sleep suggests that the corresponding emergent cortical patterns may depend on the complex interplay of the multiple brain regions controlling sleep regulation, and on the coupling of different brain rhythms [59, 60, 61, 62, 63, 64, 65, 66]. This further confirms that the AAD and the Omori law are related to the alpha rhythm dominating the awake resting-state, and suggests distinct generative mechanisms for the cascading process during NREM sleep. Despite the here-reported difference in avalanche dynamics between sleep and awake restingstate, neuronal avalanches during sleep show power-law size and duration distributions consistent with criticality [67, 68], as observed in the awake resting-state [36, 37].

We have shown that the distribution of quiet times between consecutive neural cascades, *P*(Δ*t*), exhibits two distinct power-law regimes, and related this behavior to the presence of dominant alpha oscillations. The quiet time distribution was previously investigated in other systems without alpha oscillations, and its behavior conditioned to the minimum avalanche size—i.e., *P*(Δ*t*|*s > s*_*c*_), with *s* the avalanche size and *s*_*c*_ a threshold value—was also studied [39, 35, 49]. In cortex slice cultures with up and down-states and nested theta-gamma oscillations, *P*(Δ*t*) was found to follow a non-monotonic behavior with a power-law followed by a hump and a faster decay, the hump being located at the characteristic time of the slow up/down-state oscillations [52, 39, 40]. Furthermore, it was shown that, due to the presence of up- and down-states, the conditional distributions *P*(Δ*t*|*s > s*_*c*_) do not collapse onto a unique scaling function when quiet times are rescaled by the mean quiet time, ⟨Δ*t*⟩. Most importantly, and in line with our findings, *P*(Δ*t*|*s > s*_*c*_) at the characteristic time of the theta oscillations (200 ms) was found to be independent of *s*_*c*_ [39]. On the other hand, in freely behaving rats a double power-law scaling function was found to describe the *P*(Δ*t*|*s > s*_*c*_) for a range of *s*_*c*_ [35]. In contrast with these findings and ours (Figs. S4-S5), the same analysis in zebrafish, where no oscillations are present, has shown that the quiet time distributions for different thresholds *s*_*c*_ collapse onto a unique scaling function that is well described by a Gamma distribution [49]. Overall, these observations indicate that the presence of prominent neural oscillations is connected with non-homogeneous forms of the quiet time distribution, as well as a peculiar relationship between Δ*t* and avalanche sizes *s*, where the transition between distinct scaling behaviors coincides with the characteristic period of the dominant oscillations.

Our analysis of neural activity cascades in relation to brain rhythms lays the basis for a unifying view of two complementary approaches to neural synchronization—neuronal avalanches and oscillations. On the one hand, brain rhythms have characteristic times and amplitudes, and are, by definition, a property of the integrated electromagnetic signals arising from the superposition of synaptic currents from large neural populations. On the other hand, neuronal avalanches exhibit scale-free, power-law statistical properties, and show a consistent spatio-temporal organization in terms of discrete events, from sequences of spiking neurons [69, 70, 49, 71, 72] to clusters of extreme amplitude fluctuations in LFP, EEG, and MEG sensor arrays [33, 34, 36, 37]. The coexistence of neuronal avalanches and oscillations was previously investigated in mature cortex slice cultures, in rodents, and in non-human primate, where nested *θ/β/γ*–oscillations embedded in avalanches were reported [38, 39, 41], and a hierarchical organization of *θ* and *γ* oscillations was identified [39]. In line with our observations pointing to substantial differences between cascading processes during distinct physiological states—i.e. awake resting-state and NREM sleep—, studies of spontaneous activity in mature cortex slice cultures showed that avalanche dynamics are highly sensitive to the excitation-inhibition balance [52, 53, 54]. Furthermore, a temporal structure of neuronal avalanches consistent with the Omori law was also identified in [55]. Recently, the relation between neuronal avalanches, *γ* oscillations, and emergent signatures of critical dynamics has been studied in non-human primates [41]. On the other hand, in the human brain, the relationship between avalanche dynamics and oscillations had not been scrutinized to date. Indeed, simultaneous investigations of oscillations and avalanches in the human brain selectively focused on long-range temporal correlations in alpha amplitude fluctuations and on avalanche scaling features [17, 73]. At the same time, models showing simultaneous emergence of avalanches and oscillatory behaviors mostly concentrated on underlying mechanisms or signatures of criticality [22, 74, 75, 76, 77, 23]. In particular, a quantitative analysis of awake resting-state brain activity through a class of adaptive neural networks recently linked the coexistence of alpha oscillations and avalanches to proximity to a non-equilibrium critical point at the onset of self-sustained oscillations [23, 78]. In this context, the present study establishes the first functional and dynamical links between neural oscillations and avalanches in the awake resting-state, uncovering a deep relationship between two collective phenomena with antithetic features—scale-free avalanches and scale-specific brain rhythms.

Overall, the analysis of accessible, near-synchronous collective behaviors shows that the alpha rhythm functions as a pacemaker for network excitability during the awake resting-state. The here-identified AAD correlates with alpha rhythmicity and shapes neural activity on multiple timescales, from a few hundreds of milliseconds to several seconds, indicating that alpha regulates the timing of inhibition and excitation bouts in the awake resting-state brain activity. The results suggest a unifying view of the pulsed inhibition function and the “waxing and waning” phenomenon, where the latter is a mechanism that regulates long-term, resting wake network excitability. Future work will focus on the role of AAD in information processing. In this respect, the approach we put forward will allow to (i) directly verify, and potentially extend, the pulsed inhibition hypothesis [30, 8] through the analysis of the AAD in relation to processing of sensory stimuli, and (ii) clarify the functional role of the “waxing and waning” phenomena. More generally, our approach outlines a coherent view for the dichotomy of scale-specific oscillations and scale-free avalanches [38, 39, 54, 41, 23], and demonstrates a functional and informative connection between these two phenomena. This may prove useful to dissect collective neural dynamics underlying brain oscillations in all those contexts where simultaneous recordings of single-cells and coarse-grained signals are out of reach, harvesting information from the analysis of neural cascading processes that would not be accessible otherwise.

### Limitations of the study

We identified hallmarks of alpha-mediated pulses of attenuation and amplification of neural activity cascades from MEG and EEG recordings. However, simultaneous multi-scale recordings will be needed to relate the reported large-scale dynamics with the collective behavior of local neural population, in particular modulation of inhibitory versus excitatory neural populations activity. This is a key step towards assessing whether alpha oscillations drive alternating pulses of inhibition and excitation, as our results seems to indicate. To further validate the link between AAD and alpha rhythm, a future analysis of alpha-depressed restingstate will be needed—to be also compared with the reported evidence of AAD absence during NREM sleep.

In addition, we note that the alpha rhythm is also present during REM sleep [4, 79]. However, here, we limited our investigation to the functional role of alpha oscillations in resting wake brain activity because we were interested in finding evidence of alpha-mediated pulsed inhibition [30]. The presence of AAD in relation to alpha oscillations during REM sleep should be of great interest for future work.

## STAR METHODS

## RESOURCE AVAILABILITY

### Lead contact

Further information and requests for resources and reagents should be directed to and will be fulfilled by the lead contact, Fabrizio Lombardi.

### Materials availability

This study did not generate new unique reagents.

### Data and code availability

- This study did not generate any new datasets. Data reported in this paper will be shared by the lead contact upon request.
- This study does not report original code.
- Any additional information required to reanalyze the data reported in this paper is available from the lead contact upon request.

## EXPERIMENTAL MODEL AND STUDY PARTICIPANT DETAILS

### MEG of the eyes-closed awake resting-state

This study uses data sets previously collected at the NIMH. The 4-min MEG data are the same as used in [36]. The 40-min data are the same as used in [80]. Ongoing brain activity was recorded from 100 healthy participants (38 males and 66 females; age, 31.8 ± 11.8 y) in the MEG core facility at the NIMH (Bethesda, MD, USA) for a duration of 4 min (eyes closed), and from three healthy female (age range 24–29) participants for a duration of 40 min (eyes closed). All experiments were carried out in accordance with NIH guidelines for human subjects. For the present studies 73 subjects [70 (4 min) + 3 (40 min)] with a dominant alpha peak and AAD transition were selected.

### EEG of the eyes-closed awake resting-state

Resting-state EEG was recorded for 3 min (eyes closed) from six right-handed healthy volunteers (age range 22– 27). Participants had no history of neurological or psychiatric diseases and had normal or corrected-to-normal vision. All participants gave written informed consent, and were paid for their participation. The study was approved by a local ethics committee (Ben-Gurion University) and was in accordance with the ethical standards of the Declaration of Helsinki.

### Sleep EEG

The data analyzed in this study were extracted from overnight polysomnography (PSG) recordings acquired at the Parma Sleep Disorders Centre. Ten healthy subjects, 5 males and 5 females (age range 28–53 y; average age was 39,6 y) were selected after an entrance investigation based on the following inclusion criteria: (i) absence of any psychiatric, medical, and neurological disorders; (ii) normal sleep/wake habits without any difficulties in falling or remaining asleep at night. A personal interview integrated by a structured questionnaire confirmed good vigilance level during the daytime; and (III) no drug intake at the time of the PSG or the month before.

## METHOD DETAILS

### Data acquisition and pre-processing

### MEG of the eyes-closed awake resting-state

The sampling rate was 600 Hz, and the data were band-pass filtered between 1 and 80 Hz. Power-line interferences were removed using a 60-Hz notch filter designed in Matlab (Mathworks). The sensor array consisted of 275 axial first-order gradiometers. Two dysfunctional sensors were removed, leaving 273 sensors in the analysis. Analysis was performed directly on the axial gradiometer waveforms.

### EEG of the eyes-closed awake resting-state

EEG was recorded using the g.tec HIamp system (g.tec, Austria) with 64 gel-based electrodes (AgCl electrolyte gel). Electrodes were positioned according to the standard 10/20 system with linked ears reference. Impedances of all electrodes were kept below 5 kΩ. Data were pre-processed using a combination of the EEGLAB Matlab toolbox [81] routines and custom code. After high-pass filtering (cut-off 1 Hz), a customized adaptive filter was applied to suppress line-noise. This was followed by Artifact Subspace Reconstruction [82], re-referencing to the mean, and low-pass filtering (cutoff 60 Hz). Subsequently, an ICA (independent component analysis) algorithm was applied to the data [83]. The resulting ICs were evaluated automatically for artifacts by combining spatial, spectral and temporal analysis of ICs. ICs identified as containing ocular, muscular, or cardiac artifacts were removed from the data.

### Sleep EEG

Full-night unattended PSG recordings were performed with EOG (2 channels), EEG (19 channels in 7 subjects, Ag/AgCl electrodes placed according to the 10–20 International System referred to linked-ear lobes: Fp2, F4, C4, P4, O2, F8, T4, T6, Fz, Cz, Pz, Fp1, F3, C3, P3, O1, F7, T3, T5; 25 channels in 3 subjects: CP3, CP4, C5, C6, C2, C1, FC4, FC3, F4, C4, P4, O2, F8, T4, T6, Fz, Cz, Pz, F3, C3, P3, O1, F7, T3, T5), EMG of the submentalis muscle, ECG, body position monitor, and signal for SpO2 (pulse-oximetry O2-saturation). Sleep EEG recordings were obtained using a Brain Quick Micromed System 98 (Micromed, SPA) recording machine. The institutional Ethical Committee Area Vasta Emilia Nord approved the study (protocol no. 19750). Sleep was scored visually in 30-s epochs using standard criteria [4].

### Analysis of collective neural activity

*MEG and EEG of the eyes-closed awake resting-state*.. For each MEG (EEG) sensor, positive and negative deflections in the MEG (EEG) signal were detected by applying a threshold *h* at ± *n*SD. Comparison of the signal distribution to the best fit Gaussian indicates that the two distributions start to deviate from one another around 2.7SD [36]. Thus, thresholds smaller than 2.7SD will lead to the detection of many events related to noise in addition to real events, whereas much larger thresholds will miss many of the real events. To avoid noise-related events while preserving most of relevant events, in this study, we used threshold values *h >* 2.9SD. To ensure a similar event rate across different sets of recordings (4-min MEG, 40-min MEG, 3-min EEG), we used the following *h* values: 3.3 SD for 4-min MEG; 3 SD for 40-min MEG, and 3 SD for 3-min EEG. In each excursion beyond the threshold, a single event was identified at the most extreme value (maximum for positive excursions and minimum for negative excursions). Data were binned using a time window *ϵ* = 2*T*_*meg*_ = 3.3 ms and *ϵ* = *T*_*eeg*_ = 4 ms for MEG and EEG data, respectively. *T*_*meg*_ = 1.67 ms was the sampling interval for MEG recordings, while *T*_*eeg*_ = 4 ms was the sampling interval for EEG recordings. A neural activity cascade, or avalanche, was defined as a continuous sequence of time bins in which there was at least one event on any sensor, ending with a time bin with no events on any sensor. The size of an avalanche, *s*, was defined as the number of events in the avalanche. For more details, see [36]. The size of an avalanche can be equivalently defined as the sum over all channels of the absolute values of the signals exceeding the threshold (Fig. S10).

### Sleep EEG

To identify avalanches during sleep, positive and negative deflections were detected by applying a threshold *h* ± 2SD (comparison of the signal distribution to the best fit Gaussian indicates that the two distributions start to deviate from one another at around 2SD). An avalanche was then defined as a continuous time interval in which there was at least one EEG channel supra-threshold. Due to the reduced number of electrodes in EEG sleep recordings, the size of an avalanche was defined as the sum over all channels of the absolute values of the signals exceeding the threshold. This definition is equivalent to the definition of avalanche size as the number of electrodes with positive (negative) deflections exceeding the threshold *h* [33] (Fig. S10).

### Conditional probabilities analysis

Each recording results in a sequence of avalanches *a*_*i*_, and corresponding sizes *s*_*i*_ (Fig. 1). The quantity Δ*s* = *s*_*i*+1_ – *s*_*i*_ is the difference between the sizes of two consecutive avalanches *a*_*i*_ and *a*_*i*+1_, and is used to study their monotonic relation—i.e. whether *s*_*i*_ is more likely to be smaller or larger than *s*_*i*+1_—as a function of the quiet times Δ*t* occurring in between. To this end, the following conditional probability is defined,

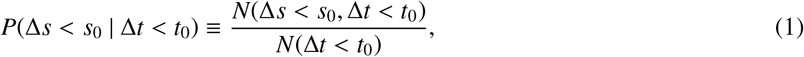

where *N*(Δ*s < s*_0_, Δ*t < t*_0_) is the number of avalanche couples which are separated by a quiet time Δ*t* shorter than a given *t*_0_ and whose size difference Δ*s* is smaller than a given *s*_0_, and *N*(Δ*t < t*_0_) is the number of quiet times Δ*t* shorter than *t*_0_. *P*(Δ*s < s*_0_|Δ*t < t*_0_) gives the probability that two consecutive avalanches separated by a Δ*t* shorter than *t*_0_ are such that Δ*s < s*_0_, with *s*_0_ a positive or negative integer. The conditional probability defined by Eq.(1) assesses the monotonic relation between consecutive avalanches as a function of the quiet time separating them, thus, providing a detailed picture of the correlation landscape of the avalanche process. To measure the strength and significance of such a relationship, for each given couple of thresholds *s*_0_ and *t*_0_, *P*(Δ*s < s*_0_|Δ*t < t*_0_) is systematically compared with the same conditional probability evaluated over 10^4^ independent surrogate avalanche time series. Surrogates are obtained by randomly reshuffling avalanche sizes while keeping fixed their starting and ending times. *P*(Δ*s < s*_0_ | Δ*t < t*_0_) are then compared with the average surrogate conditional probabilities, *P*^∗^(Δ*s < s*_0_|Δ*t < t*_0_), by analyzing the quantity

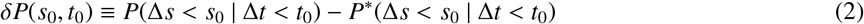

as a function of *s*_0_ for different fixed values of *t*_0_ (Fig. 3). Being *σ*^∗^(*s*_0_, *t*_0_) the standard deviation of the surrogate conditional probability *P*^∗^(Δ*s < s*_0_|Δ*t < t*_0_), if *δP*(*s*_0_, *t*_0_)|*>* 2*σ*^∗^(*s*_0_, *t*_0_), one can conclude that *P* and *P*^∗^ are significantly different (0.05 significance level), and that significant correlations exist between Δ*s < s*_0_ and Δ*t < t*_0_.

Two cases must be distinguished: *δP*(*s*_0_, *t*_0_) *>* 0 and *δP*(*s*_0_, *t*_0_) *<* 0. If *δP*(*s*_0_, *t*_0_) *>* 0, the number of avalanche couples *N*(Δ*s < s*_0_, Δ*t < t*_0_) satisfying both conditions is significantly larger in the original data than in the surrogates; namely, it is more likely to find couples satisfying both conditions in the original rather than in the surrogate avalanche time series. Hence, Δ*s* and Δ*t* are positively correlated. On the contrary, if *δP*(*s*_0_, *t*_0_) *<* 0, the number of couples *N*(Δ*s < s*_0_, Δ*t < t*_0_) satisfying both conditions is significantly larger in the surrogates than in the original data; namely, it is more likely to find couples satisfying both conditions in the uncorrelated surrogates rather than in the real avalanche time series. In this case, one says that Δ*s* and Δ*t* are negatively correlated.

### Omori law

The number of avalanches per unit time, *N*(*t*), occurring after a time *t* has elapsed from the main avalanche *A*^∗^ is computed using a time window *δt* that increases logarithmically. Denoting a set of window boundaries as *W* = (*w*_1_, *w*_2_, …, *w*_*k*_) and fixing *w*_1_ = 10 ms, the logarithmic windows fulfill the relation *w*_*i*+1_ = *w*_*i*_ · 10^*c*^, which implies that the window size is constant in logarithmic scale, i.e. *logw*_*i*+1_ – *logw*_*i*_ = *c*. The following window sizes of *c* have been used in this study: *c* = 0.1 for the *N*(*t*) of the awake resting-state (Fig. 3, Fig. S9); *c* = 0.11 for the *N*(*t*) during sleep (Fig. 4).

### Spearman’s correlation coefficient

Given two variables *X* and *Y*, the Spearman’s correlation coefficient is defined as

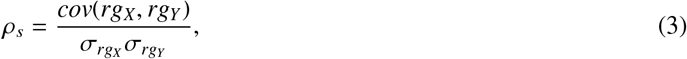

where *rg*_*X*_ and *rg*_*Y*_ are the tied rankings of *X* and *Y*, respectively, 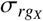 and 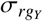 their standard deviations, and *cov*(*rg*_*X*_, *rg*_*Y*_) indicates the covariance between *rg*_*X*_ and *rg*_*Y*_.

### Surrogate signals

Surrogate signals are obtained by random phase shuffling of the original continuous signals. A Fourier transform of each sensor signal is performed, and the corresponding phases are randomized while amplitudes are preserved. The surrogate signals are then obtained by performing an inverse Fourier transform. The random phase shuffling destroys phase synchronization across cortical sites while preserving the linear properties of the original signals, such as power spectral density and two-point correlations [84]. Surrogate signals were used to generate surrogate data for Fig. 2, 4, and 5.

### Surrogate time series for correlations between Δs and Δt

To test significance of correlations between consecutive Δ*s* and Δ*t*, a surrogate sequence of avalanche sizes was generated for each subject by randomly reshuffling the original order of avalanche sizes. The Spearman’s correlation coefficient *ρ*(Δ*s*, Δ*t*) between consecutive Δ*s* and Δ*t* was calculated for each surrogate. The average Spearman’s correlation coefficient obtained from all surrogates was then compared with the average correlation coefficient calculated from the original sequences of avalanche sizes and quiet times (Fig. 1 and 4).

## QUANTIFICATION AND STATISTICAL ANALYSIS

Power law exponents were estimated using a maximum likelihood estimator [85, 44, 45]. The power law fit was compared to an exponential fit by evaluating the log-likelihood ratio *R* = *lnL*_*p*_*/L*_*e*_, where *L*_*p,e*_ = _*i*=1_ *p*_*p,e*_(*x*_*i*_) is the likelihood. *R* is positive if the data are more likely to follow a power law distribution, and negative if the data are more likely to follow exponential distribution. The statistical significance for *R* was estimated as in [44, 85]. Following [85], the *p*-value associated to *R* is given by

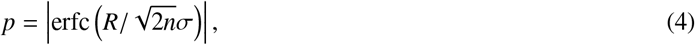

where *σ*^2^ is the variance of the data [85] and

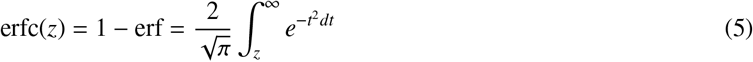

is the complementary Gaussian error function [85].

Power-law exponent and fitting parameters for quiet time distributions, *P*(Δ*t*), for the autocorrelation *C*(*t*), and for the function *N*(*t*) were evaluated on pooled data, unless otherwise stated (Fig. 1, 3, and 4). The corresponding error reported in the main text and in the figure caption is the error on the fit, unless otherwise stated.

Within each data group (MEG, EEG resting wake, EEG sleep), the Spearman’s correlation coefficients *ρ*(Δ*s*, Δ*t*) were evaluated for each subject. The values reported in the main text and in the figure captions are (mean ± SD) (Fig. 1 and 4). Significance of correlations was assessed comparing the average correlation coefficient calculated from the original sequences of avalanche sizes and quiet times with the correlation coefficient calculated from surrogate time series (Fig. 1 and 4). Pairwise comparisons were conducted using two-tailed Student’s *t*-test.

## Supplemental information

Supplemental information can be found online at.

### Acknowledgements

This research was funded in whole, or in part, by the Austrian Science Fund (FWF) (grant no. PT1013M03318 to F.L.). For the purpose of open access, the author has applied a CC BY public copyright licence to any Author Accepted Manuscript version arising from this submission. The study was supported by the European Union Horizon 2020 research and innovation program under the Marie Sklodowska-Curie action (grant agreement No. 754411 to F.L.), and in part by the NextGenerationEU through the grant STARS@UNIPD to F.L. (project “Brain criticality and information processing (BRAINCIP)”). LdA acknowledges support from the Italian MIUR project PRIN2017WZFTZP and partial support by NEXTGENERATIONEU (NGEU) funded by the Ministry of University and Research (MUR), National Recovery and Resilience Plan (NRRP), project MNESYS (PE0000006) — A Multiscale integrated approach to the study of the nervous system in health and disease (DN. 1553 11.10.2022). OS acknowledges support from the Israel Science Foundation, Grant No. 504/17. The work was supported in part by DIRP ZIAMH02797 to DP.

## Author contributions

Conceptualization, F.L., L.d.A., and O.S.; Methodology, F.L., L.d.A., and O.S.; Formal Analysis, F.L. and S.S.; Investigation, all authors; Writing-Original Draft, F.L.; Writing-Review & Editing, all authors; Visualization, F.L.

## Declaration of interests

The authors declare no competing interests.

## Inclusion and diversity

We support inclusive, diverse, and equitable conduct of research.

## Notes

### Competing Interest Statement

The authors have declared no competing interest.

### Summary of Updates

Title revised; Abstract revised; New Fig. 1; New supplementary figures; Fig. 2 revised. Discussion revised.

