## Supplementary Information for "Beyond pulsed inhibition: Alpha oscillations modulate attenuation and amplification of neural activity in the awake resting-state"

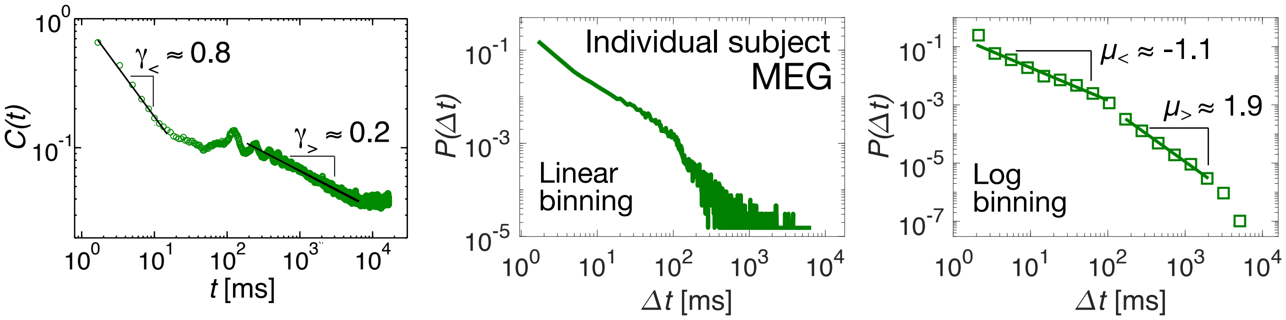


Fig. S1: **Correspondence between the power-law decays of the activity autocorrelation and the power-laws in the quiet time distribution. Related to Figure 2.** The activity auto-correlation, $C(t)$, shows two distinct power law decays (left): one at $t < 100$ ms, with $\gamma_< = 0.76 \pm 0.08$; the other one at $t > 100$ ms: $\gamma_> = 0.23 \pm 0.09$. Such power law decays are separated by a plateau region in the vicinity of $t = 100$ ms. In close correspondence with $C(t)$, the quiet time distribution (middle, linear binning; right logarithmic binning) exhibits two distinct power law regimes: $\Delta t < 100$ ms, $\mu_< = 1.12 \pm 0.08$; $\Delta t > 100$ ms, $\mu_< = 1.91 \pm 0.07$. The exponents $\gamma_<$ ($\gamma_>$) and $\mu_<$ ($\mu_>$) approximately obey the theoretical relationship $\mu_< = 2 - \gamma_<$ ($\mu_> = 2 - \gamma_>$) [1, 2, 3]. All data are from an individual 40-min MEG recording.

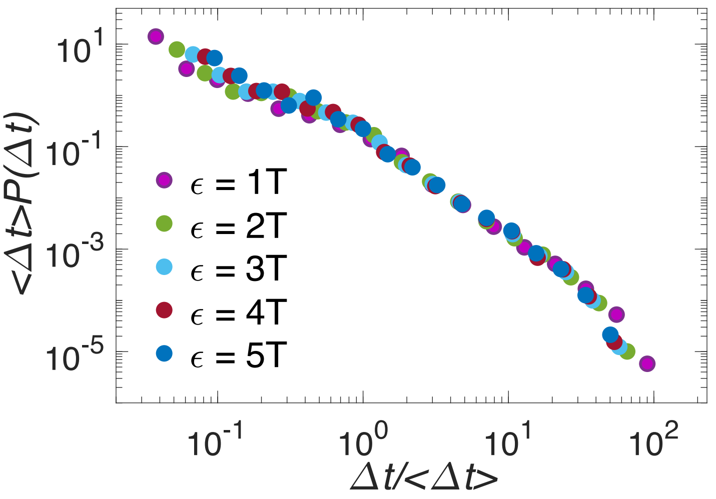


Fig. S2: **The distribution of quiet time is independent of the bin size epsilon used to define avalanches. Related to Figure 2.** Quiet time distribution for an individual 40-min MEG subject and different bin sizes $\epsilon$. Distributions are rescaled by the mean $\Delta t$, $\langle \Delta t \rangle$.

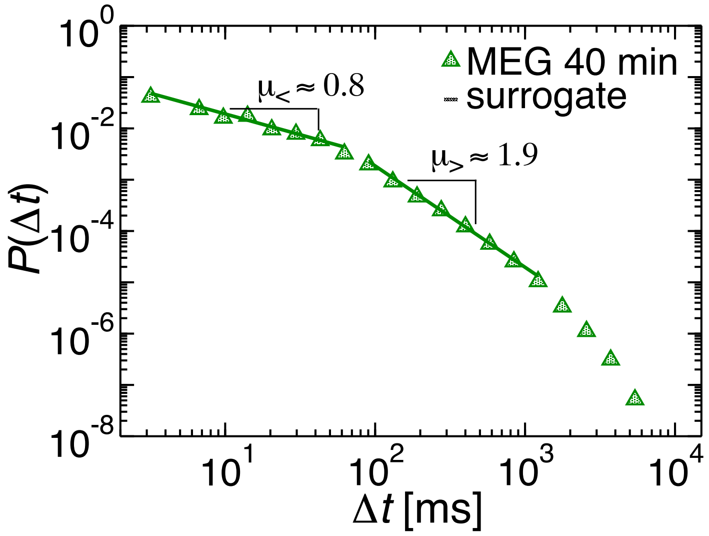


Fig. S3: **The distribution of quiet time in 40-min MEG exhibits a double power-law behavior as in 4-min MEG. Related to Figure 2.** The regime ($A_<$) for short $\Delta t$'s ($\Delta t < 100$ ms) is characterized by an exponent $\mu_< = 0.85 \pm 0.04$ (power-law versus exponential comparison: $R = 321$; $p = 0.0068$). For longer $\Delta t$'s (100 ms $< \Delta t <$ 1200 ms, regime ($A_>$)), the power law exponent is $\mu_> = 1.9160 \pm 0.0471$ (power-law versus exponential comparison: $R = 707$, $p = 3 \cdot 10^{-52}$) . The transition region between regime ($A_<$) and ($A_>$) is located around 100 ms, as in 4-min MEG and EEG data (Fig. 2).

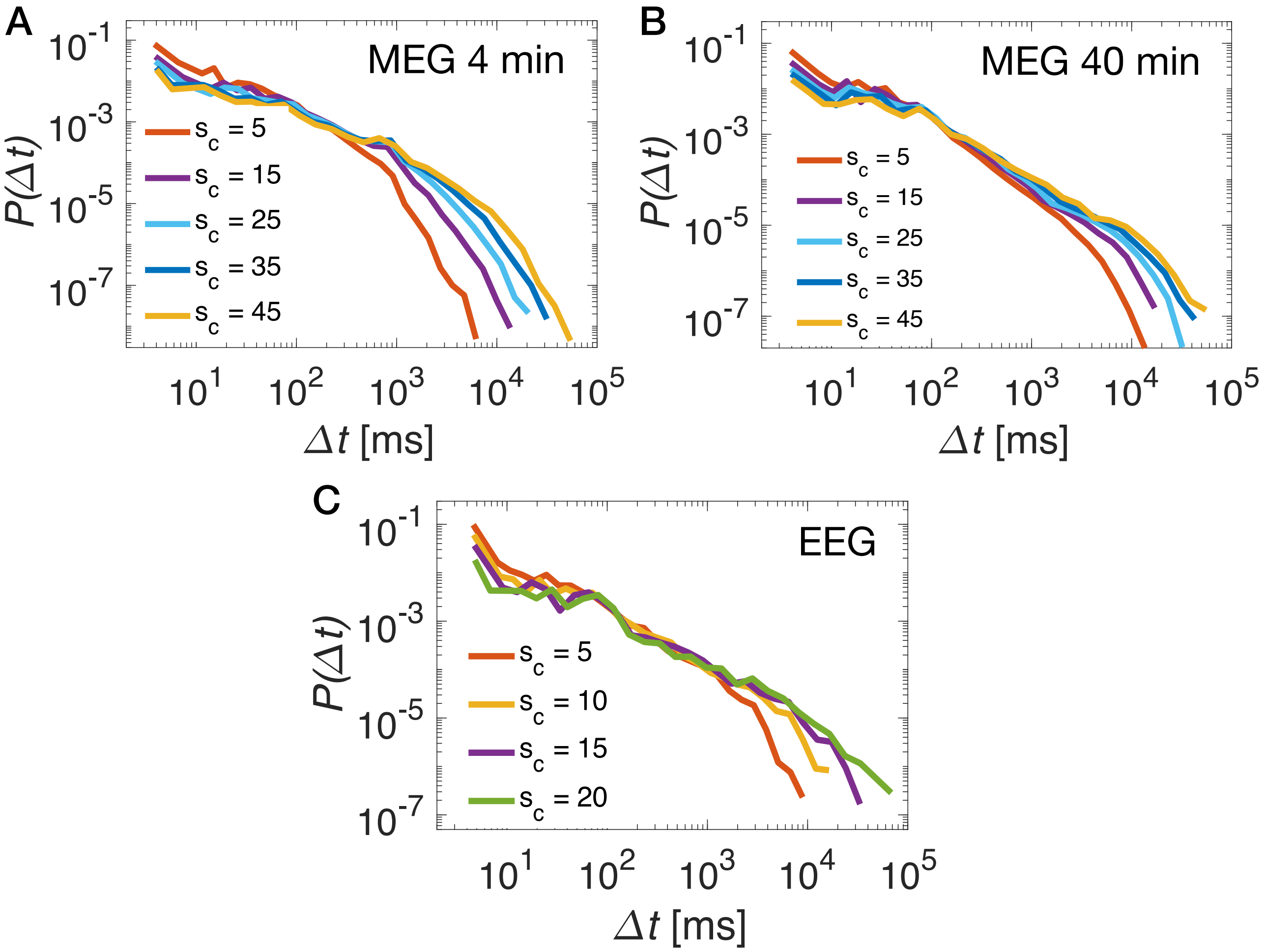


Fig. S4: **Distributions of quiet times for different minimum thresholds $s_c$ on avalanche sizes in the awake resting state. Related to Figure 2.** For increasing $s_c$ values, we observe that (i) $P(\Delta t)$ decreases for $\Delta t < 100$ ms, and tends to a plateau regime ; (ii) $P(\Delta t)$ increases for $\Delta t > 100$ ms, and shows a power-law with decreasing exponent; (iii) $P(\Delta t)$ stays approximately constant around $\Delta t = 100$ ms, i.e. $\Delta t = 100$ ms is a sort of fix point for $P(\Delta t|s_c)$.

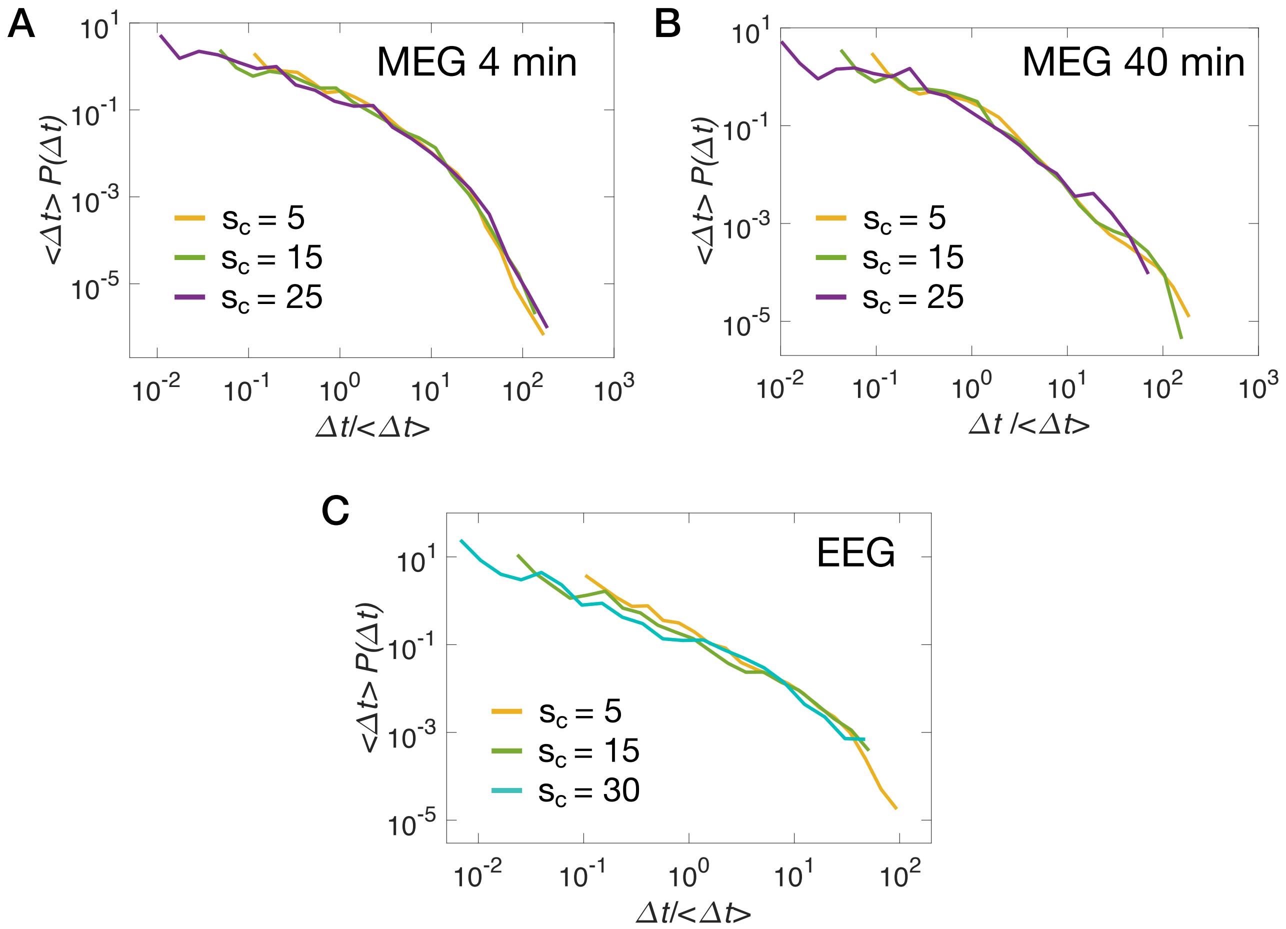


Fig. S5: **Distributions of rescaled quiet times for different minimum thresholds $s_c$ on avalanche sizes in the awake resting state. Related to Figure 2.** Distributions do not collapse onto a single curve. This is particularly evident in (B) and (C).

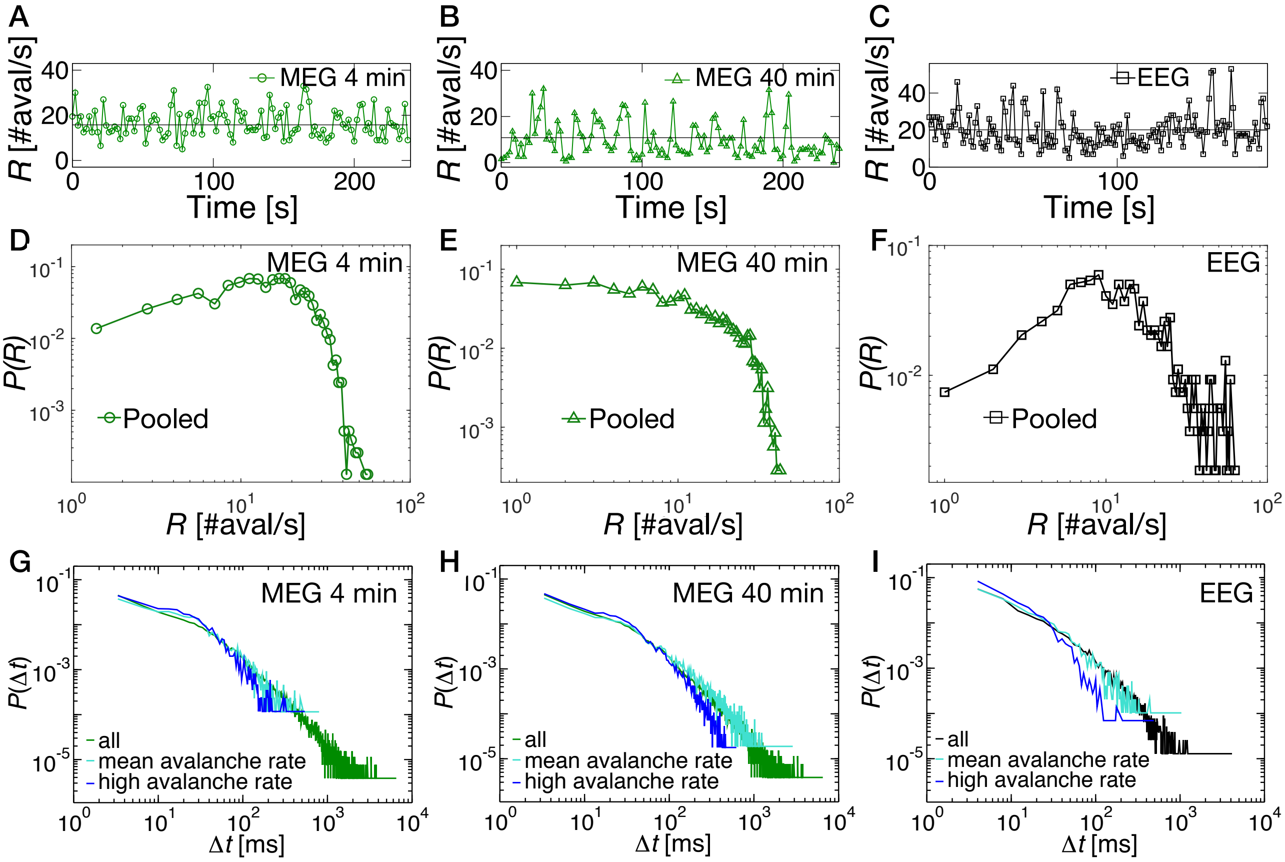


Fig. S6: **Avalanche rate analysis in the awake resting state. Related to Figure 2. (A-C)** Avalanche rate for an individual subject from the 4-min MEG dataset (A), the 40-min MEG dataset (B), and the 3-min EEG dataset (C) (thick black line = mean). Unlike the typical behavior observed in earthquake sequences (as those shown in [4]), $R(t)$ shows continuous fluctuations around the mean in all cases (slow periodic fluctuations are also present in some cases, e.g. in B). We do observe peaks in $R(t)$, but these are usually of the order of 1-2 SD, whereas for earthquakes rate jumps are as large as several order of magnitude. Thus, in the case of neuronal avalanches, the distinction between homogeneous and non-homogeneous periods may hardly apply. **(D-F)** Distribution of avalanche rate, $P(R)$, in 4-min MEG recordings (D), 40-min MEG recordings (E), and 3-min EEG recordings (F) (pooled data). In all cases, the distribution of avalanche rate, $P(R)$, does not follow a double power-law behavior, which, on the contrary, was reported for earthquakes in [4]. We observe that $P(R)$ is approximately constant with an exponential cutoff in both 4-min and 40-min MEG recordings, while for EEG data $P(R)$ tends to be peaked around a particular $R$ value. $R$ and $P(R)$ were calculated using a 1 s time window—about 100 times the most likely avalanche duration and 10 times the longest duration. This choice ensures a good trade-off between capturing the temporal fluctuations of the avalanche process and having a significant "density" of events in each window. However, the distribution $P(R)$ does not crucially depend on this choice. **(G-I)** Distribution of quiet times, $P(\Delta t)$, in periods with the avalanche rate comprised between (mean - SD) and (mean + SD) (cyan curve, mean-rate periods), and periods with avalanche rate larger than (mean + SD) (blue curve, high-rate periods). In high-rate periods, the probability of longer $\Delta t$ slightly decreases, while the probability of shorter $\Delta t$ increases, as expected. However, we observe that the general form of the quiet time distribution does not change significantly. In particular, as expected, we do not see changes at short $\Delta t$ (high rate). Moreover, we continue to observe the transition at $\approx 100$ ms. The observed differences on the tail of the distribution (low rate) should be ascribed to a lack of statistics (we are removing long $\Delta t$, which are already rare) rather than an intrinsic difference in the dynamics. This is particularly evident when looking at the quiet time distribution for the EEG data, where we have only 6 subjects recorded for 3 min. Importantly, results do not qualitatively change for higher threshold. However, the statistics becomes rather poor, and the distributions noisier.

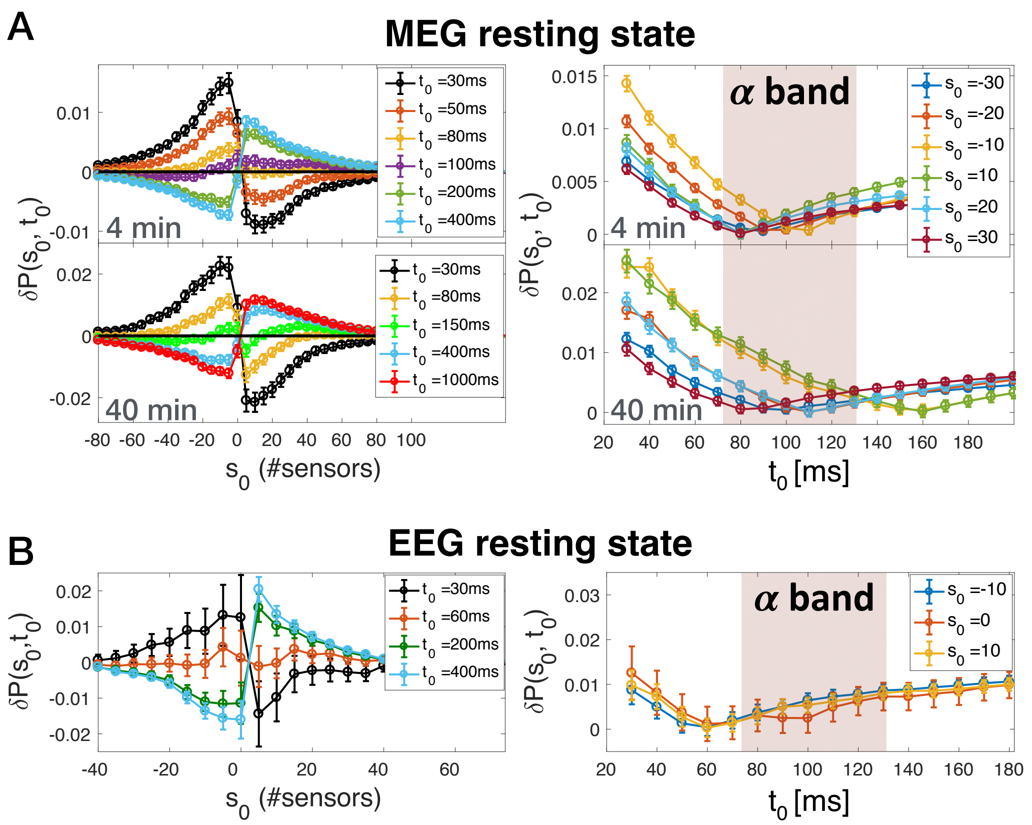


Fig. S7: **The transition from the attenuation to the amplification regime in the resting-state cascading process occurs at the characteristic time of the alpha rhythm. Related to Figure 3.** **(A)** (Left panel) $\delta P(s_0, t_0)$ as a function of the threshold $s_0$ on $\Delta s$ for different values of the threshold $t_0$ on the quiet times $\Delta t$'s for MEG resting state recordings. The error bar on each data point is two times the standard deviation $\sigma^*$ associated with the surrogates $P(s_0, t_0)^*$ (see Fig. 3 and STAR Methods). For a given threshold $t_0$, the maximum in $\delta P(s_0, t_0)$ indicates the preferred relation between consecutive avalanches separated by quiet times shorter than $t_0$. We notice that, both in the 4-min (upper panel, n = 70) and 40-min MEG recordings (lower panel, n = 3), for $t_0 < 100$ ms, $\delta P(s_0, t_0)$ has a maximum at $s_0 < 0$, and therefore an avalanche tends to be smaller than its preceding one ($s_{i+1} < s_i$, attenuation regime). On the other hand, for $t_0 > 100$ ms the maximum moves towards positive $s_0$, implying that a given avalanche tends to be larger than the preceding one ($s_{i+1} > s_i$, amplification regime). Remarkably, for $t_0 \simeq 100$ ms, $\delta P(s_0, t_0)$ is very close to zero for each $s_0$, with values that are not significantly different among each other. This indicates that the conditional probabilities evaluated on the original data are very close to those calculated from the surrogates. Hence, we find that at $t_0 \simeq 100$ ms there is not a preferred sign for $\Delta s$, indicating that $\Delta t = 100$ ms is a transition point from one dynamical regime to another. (Right panel) The absolute value $|\delta P(s_0, t_0)|$ as a function of $t_0$ for several fixed values of $s_0$. For each $s_0$, $|\delta P(s_0, t_0)|$ has a minimum around $t_0 = 100$ ms . More specifically, the minimum is always comprised between 80 ms and 110 ms for the 4-min recordings (upper panel), and between 80 ms and 160 ms for the 40-min recordings (lower panel). Such time scales correspond to the characteristic period of the $\alpha$ rhythm. **(B)** (Left panel) $\delta P(s_0, t_0)$ as a function of the threshold $s_0$ on $\Delta s$ for different values of the threshold $t_0$ on the quiet times $\Delta t$'s for EEG resting state recordings. The error bar on each data point is two times the standard deviation $\sigma^*$ associated with the surrogates $P(s_0, t_0)^*$ (see Fig. 3 and STAR Methods). As in MEG recordings, we observe a transition between two different dynamical regimes. For $t_0 < 60$ ms, $\delta P(s_0, t_0)$ has a maximum at $s_0 < 0$, and therefore an avalanche tends to be smaller than its preceding one ($s_{i+1} < s_i$, attenuation regime). On the other hand, for $t_0 > 60$ ms the maximum moves towards positive $s_0$, implying that a given avalanche tends to be larger than the preceding one ($s_{i+1} > s_i$, amplification regime). Remarkably, for $t_0 \simeq 60$ ms, $\delta P(s_0, t_0)$ is very close to zero for each $s_0$, with values that are not significantly different among each other. This indicates that the conditional probabilities evaluated on the original data are very close to those calculated from the surrogates. Hence, we find that at $t_0 \simeq 60$ ms there is not a preferred sign for $\Delta s$, suggesting indeed that $\Delta t = 60$ ms is a transition point from one dynamical regime to another. (Right panel) The absolute value $|\delta P(s_0, t_0)|$ as a function of $t_0$ for several fixed values of $s_0$. For each $s_0$, $|\delta P(s_0, t_0)|$ has a minimum around $t_0 = 60$ ms. The error bar on each data point is two times the standard deviation $\sigma^*$ associated with the surrogates $P(s_0, t_0)^*$ (see Fig. 3 and STAR Methods).

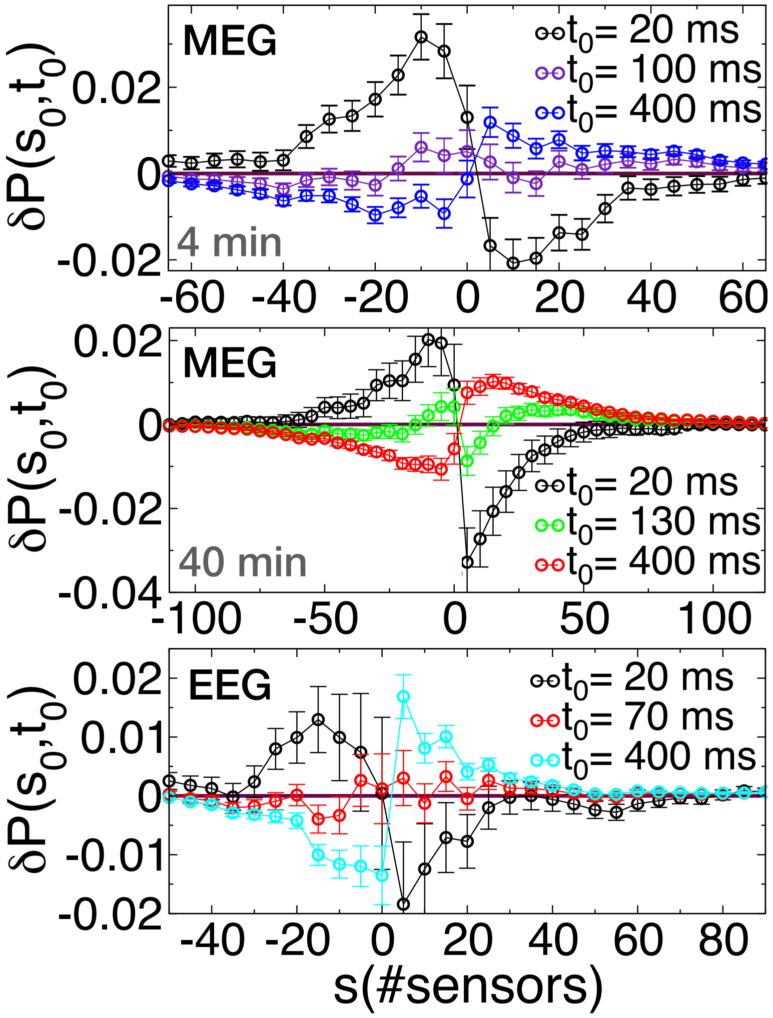


Fig. S8: **The attenuation-amplification transition in the neural cascading dynamics is robust and consistent across individual subjects. Related to Figure 3. (Top)** $\delta P(s_0, t_0)$ as a function of the threshold $s_0$ for an individual 4-min MEG recording. **(Middle)** $\delta P(s_0, t_0)$ as a function of the threshold $s_0$ for an individual 40-min MEG recording. **(Bottom)** $\delta P(s_0, t_0)$ as a function of the threshold $s_0$ for an individual EEG recording. The error bar on each data point is two times the standard deviation $\sigma^*$ associated with the surrogates $P^*(s_0, t_0)$ (Materials and Methods). For a given threshold $t_0$, the maximum in $\delta P(s_0, t_0)$ indicates the preferred relation between consecutive avalanches separated by quiet times shorter than $t_0$. In all the three cases, we notice that for $t_0 < 100$ ms $\delta P(s_0, t_0)$ has a maximum at $s_0 < 0$, and therefore an avalanche tends to be smaller than its preceding one ($s_{i+1} < s_i$, attenuation regime). On the other hand, for $t_0 > 100$ ms the maximum moves towards positive $s_0$, implying that a given avalanche tends to be larger than the preceding one ($s_{i+1} > s_i$, amplification regime). Remarkably, for $t_0 \simeq 100$ ms, $\delta P(s_0, t_0)$ is very close to zero for each $s_0$, indicating that the conditional probabilities evaluated on the original data are very close to those calculated from the surrogates. Hence, we find that at $t_0 \simeq 100$ ms there is not a preferred sign for $\Delta s$. This indicates that, both in pooled data and individual subjects, $\Delta t \approx 100$ ms is a transition point from the attenuation to the amplification regime.

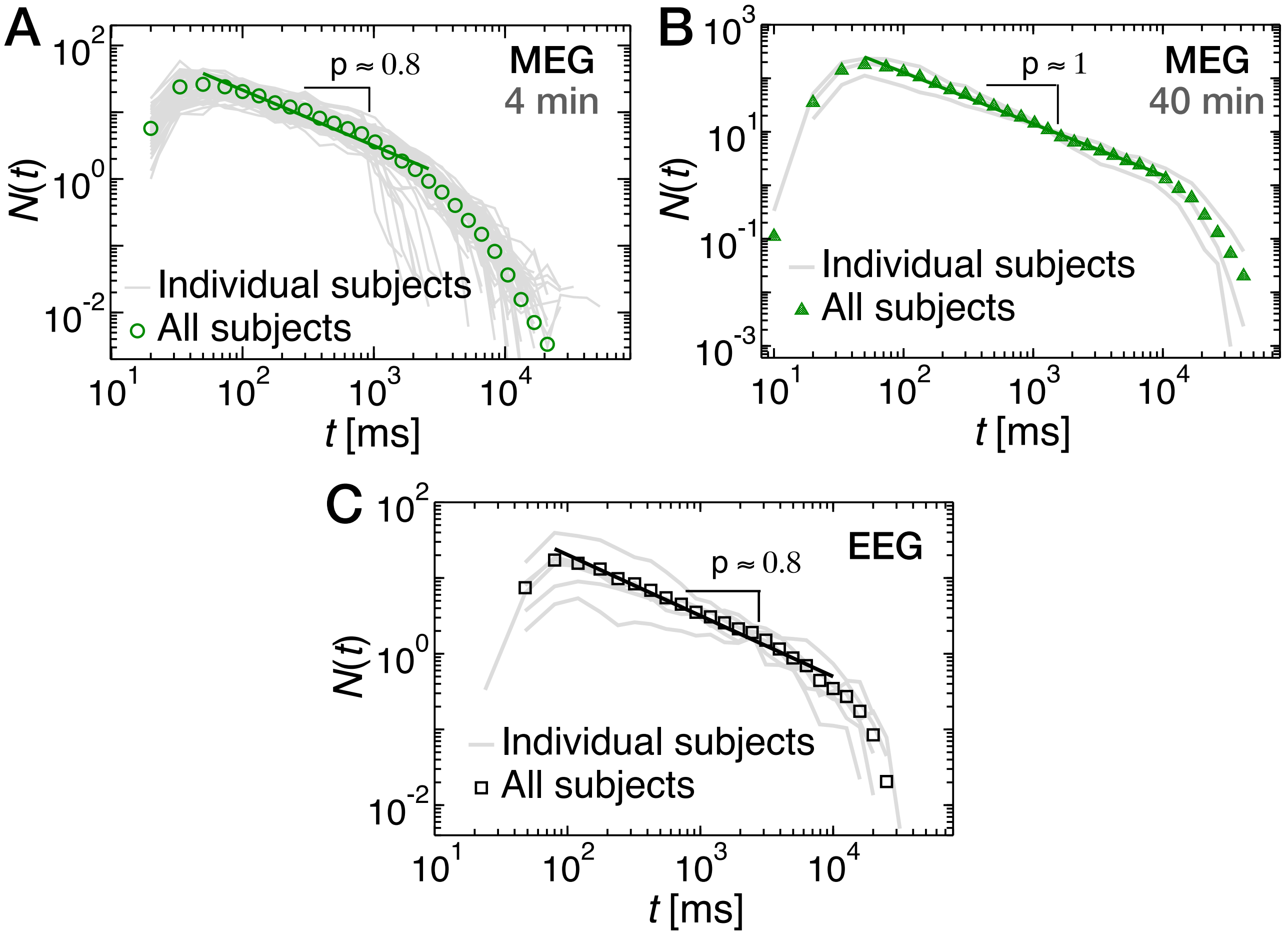


Fig. S9: **The number of avalanches per unit time, $N(t)$, occurring after a main avalanche ($A^* > 30$) obeys the Omori law. Related to Figure 4.** $N(t)$ decreases roughly as the reciprocal of the time $t$ elapsed from the main shock, i.e. $N(t) \propto t^{-p}$. **(A)** $N(t)$ for 4-min MEG recordings from individual subjects (grey lines), and pooled subjects (green circles). Power-law fit (green thick line): $p = 0.8326 \pm 0.0453$. **(B)** $N(t)$ for 40-min MEG recordings from individual subjects (grey lines), and pooled subjects (green triangles). Power-law fit (green thick line): $p = 0.9596 \pm 0.0137$. **(C)** $N(t)$ for EEG recordings from individual subjects (grey lines), and pooled subjects (black squares). Power-law fit (black thick line): $p = 0.8041 \pm 0.0277$.

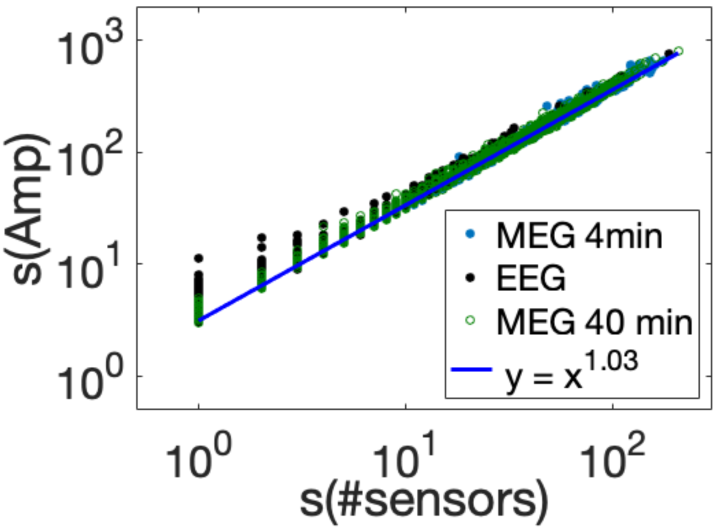


Fig. S10: **Scatter plot between avalanche sizes $s(\#sensors)$ measured as number of active sensors, and avalanche sizes $s(Amp)$ measured as the sum of the signal amplitudes across active sensors in MEG and EEG recordings of an individual subject. Related to STAR Methods and Figure 1.** We find that $s(Amp) \propto s(\#sensors)^{1.03}$. This implies that the definition of avalanche size as the number of active sensors is equivalent to the definition based on the sum of the signal amplitudes across active sensors in both MEG and EEG recordings (STAR Methods).
